# Novel sterol binding domains in bacteria

**DOI:** 10.1101/2022.05.15.491920

**Authors:** Liting Zhai, Amber C. Bonds, Clyde A. Smith, Hannah Oo, Jonathan Chiu-Chun Chou, Paula V. Welander, Laura M. K. Dassama

## Abstract

Sterol lipids are widely present in eukaryotes and play essential roles in signaling and modulating membrane fluidity. Although rare, some bacteria also produce sterols, but their function in bacteria is not known. Moreover, many more species, including pathogens and commensal microbes, acquire or modify sterols from eukaryotic hosts through poorly understood molecular mechanisms. The aerobic methanotroph *Methylococcus capsulatus* was the first bacterium shown to synthesize sterols, producing a mixture of C-4 methylated sterols that are distinct from those observed in eukaryotes. C-4 methylated sterols are synthesized in the cytosol and localized to the outer membrane, suggesting that a bacterial sterol transport machinery exists. Until now, the identity of such machinery remained a mystery. In this study, we identified three novel proteins that may be the first examples of transporters for bacterial sterol lipids. The proteins, which all belong to well-studied families of bacterial metabolite transporters, are predicted to reside in the inner membrane, periplasm, and outer membrane of *M. capsulatus,* and may work as a conduit to move modified sterols to the outer membrane. Quantitative analysis of ligand binding revealed their remarkable specificity for 4-methylsterols, and crystallographic structures coupled with docking and molecular dynamics simulations revealed the structural bases for substrate binding by two of the putative transporters. Their striking structural divergence from eukaryotic sterol transporters signals that they form a distinct sterol transport system within the bacterial domain. Finally, bioinformatics revealed the widespread presence of similar transporters in bacterial genomes, including in some pathogens that use host sterol lipids to construct their cell envelopes. The unique folds of these bacterial sterol binding proteins should now guide the discovery of other proteins that handle this essential metabolite.

## Introduction

Sterol lipids are ubiquitous and essential components of eukaryotic life, playing vital roles in intra- and intercellular signaling, stress tolerance, and maintaining cell membrane integrity (Bi and Liao, 2010; Bloch, 1991; Huang et al., 2016; Miao et al., 2002). Although sterol synthesis is often considered to be a strictly eukaryotic feature, several bacteria have been shown to also produce sterols (Bloch, 1991; Ourisson et al., 1987). Bacterial sterols were first discovered in *Methylococcus capsulatus* more than 40 years ago (BIRD et al., 1971) and initially, were only observed in a few isolated aerobic methanotrophs (γ-Proteobacteria) and a few myxobacteria (δ-Proteobacteria) (Bode et al., 2003; Kohl et al., 1983; Patt and Hanson, 1978; Schouten et al., 2000). Subsequent comparative genomics analyses have revealed the potential to produce sterol in a variety of bacterial groups including Planctomycetes, α-Proteobacteria, and Bacteroidetes (Bode et al., 2003; Pearson et al., 2003; Wei et al., 2016). Sterols produced by bacteria, however, tend to differ from eukaryotic sterols in both structure and in their biosynthetic pathways.

Sterol synthesis in both bacteria and eukaryotes requires the cyclization of the linear substrate oxidosqualene by an oxidosqualene cyclase (Osc) to generate either lanosterol or cycloartenol (Abe, 2014). However, fungi, vertebrates, and plants further modify these initial cyclization products to synthesize ergosterol, cholesterol, and stigmasterol, respectively (Desmond and Gribaldo, 2009). These biochemical transformations include demethylations, isomerizations, saturations, and desaturations that are essential for sterols to function properly in eukaryotes (Nes et al., 1993; Xu et al., 2005). Although bacterial production of fully modified sterols such as cholesterol is rare (Lee et al., 2023), some bacteria, including several aerobic methanotrophs and myxobacteria, do modify their sterols. For example, *M. capsulatus* produces sterols that are demethylated once at the C-4 and C-14 positions and contain a unique desaturation between C-8 and C-14 (Bouvier et al., 1976) (Fig.1A). In addition, studies have revealed that bacterial proteins required to modify sterols can differ from the canonical eukaryotic sterol modifying proteins. Examples are the recently identified sterol demethylase proteins, SdmA and SdmB, in *M. capsulatus* that use O_2_ to remove one methyl group at the C-4 position; the enzymes are mechanistically distinct from the eukaryotic C-4 demethylase enzymes (Gachotte et al., 1998; Lee et al., 2018; Rahier, 2011).

**Figure 1.**
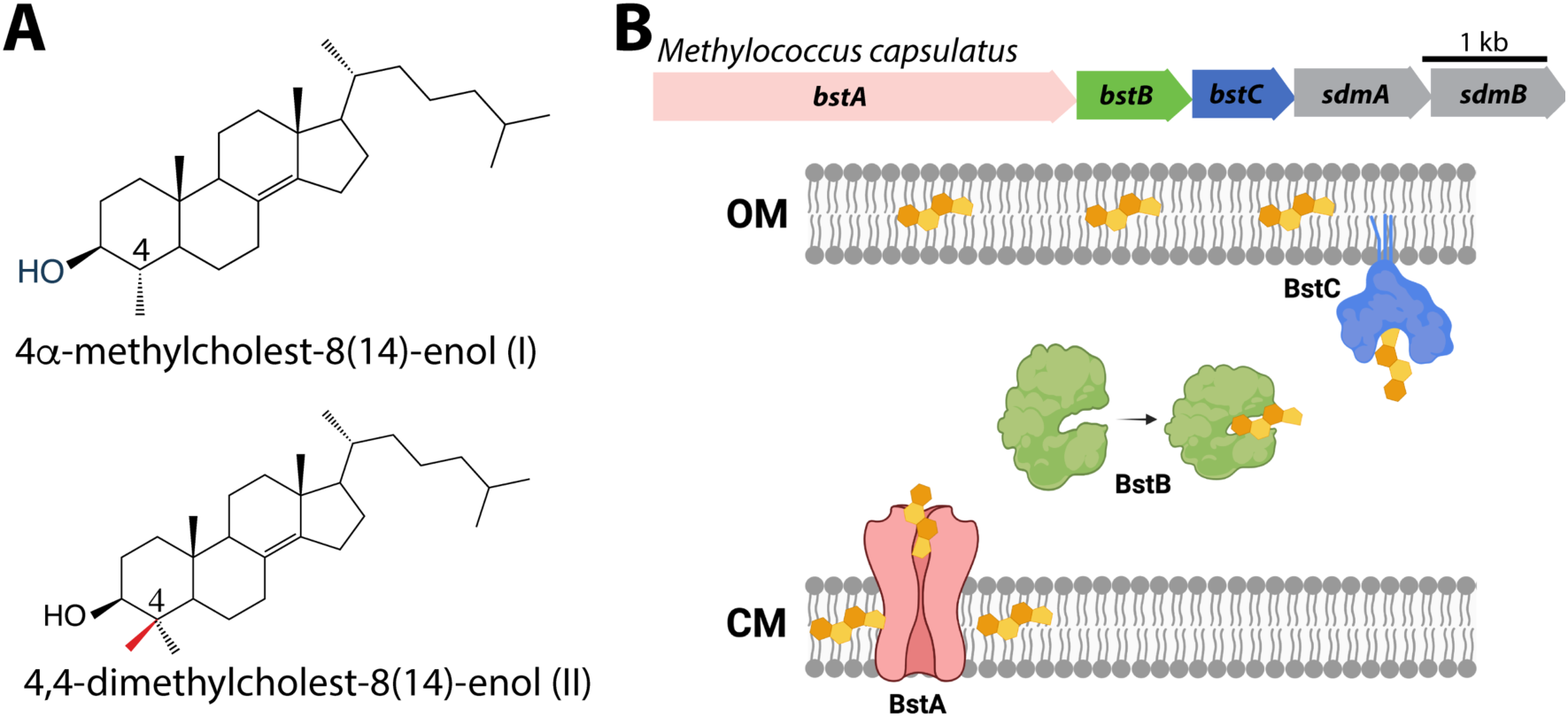
(A) Structures of C-4 methylated sterols synthesized by *Methylococcus capsulatus*. (B) Operon of sterol biosynthesis and transport genes and (top) and schematic description of their localization. OM: outer membrane; CM: cytoplasmic membrane.

Although we have a better grasp of the taxonomic distribution of sterol synthesis in bacteria and of the distinct bacterial proteins involved in their biosynthesis, the function of these lipids in bacteria remains a mystery. It has been posited that bacterial sterols play a role in modulating the fluidic properties of the cytoplasmic membrane, similar to what is observed for cholesterol and ergosterol in eukaryotic cells (Miao et al., 2002; Parks et al., 1995; Summons et al., 2006). However, studies in *M. capsulatus* have demonstrated that the distribution of sterols differs considerably from what occurs in eukaryotes. Approximately 90% of the free sterol pool in eukaryotic cells is found in the cytoplasmic membrane (Jacquier and Schneiter, 2012) while in *M. capsulatus*, 75% of sterols are localized to the outer membrane (JAHNKE et al., 1992). The outer membrane is the external asymmetric membrane of Gram-negative bacteria and differs from the cytoplasmic membrane in that its outer leaflet is composed of lipopolysaccharides (LPS) in addition to phospholipids (Ruiz et al., 2006). Additionally, sterols that differ by only a single methylation in the core structure can produce significantly different effects in terms of membrane fluidity and stability (Bacia et al., 2005). Given that *M. capsulatus* primarily produces sterols with one or two C-4 methyl groups, any sterol-membrane interactions in *M. capsulatus* could differ significantly from what is observed in eukaryotic membranes with cholesterol and ergosterol, which lack methyl groups at the C-4 position.

Interestingly, the 4,4-dimethyl sterols and 4-monomethyl sterols found in *M. capsulatus* also exist as precursors of pathway end-products in eukaryotes and accumulation of C-4 methylated sterols has been associated with a variety of eukaryotic processes. The accumulation of C-4 methylated intermediates is considered to be the cause of human genetic diseases known as sterolosis (He et al., 2011; König et al., 2000; McLarren et al., 2010) and 4,4-dimethyl sterols such as lanosterol in the brain are implicated in Parkinson’s disease (Lim et al., 2012). In a variety of yeast species, C-4 methylated intermediates can regulate cellular processes associated with hypoxia and other conditions of cellular stress (Hughes et al., 2007; Serratore et al., 2017; Todd et al., 2006). Although the function of 4,4-dimethyl sterols and 4-monomethyl sterols in bacteria is still elusive, a functional role for these specific sterols beyond maintaining membrane fluidity and integrity seems plausible.

To better understand the significance of sterol utilization in the bacterial domain, a fuller understanding of the molecular mechanisms controlling sterol production and localization is needed. Given the observed distribution of sterols in the outer membrane of *M. capsulatus*, we hypothesized that transporters specific for C-4 methylated sterols must exist that can shuttle these substrates to the outer membrane. Trafficking of sterols to various organelles in eukaryotic cells is not fully understood (Jacquier and Schneiter, 2012) and impaired sterol transport is related to a variety of defects including lysosomal storage diseases (Vance and Peake, 2011). Identifying and characterizing bacterial proteins that transport sterol could reveal novel sterol binding motifs and folds and will provide meaningful insights into protein-sterol interactions and lipid trafficking more broadly.

In this study, we present three putative bacterial sterol transport proteins in *M. capsulatus* that exhibit remarkable specificity for C-4 methylated sterols. We first used bioinformatics to identify a cytoplasmic membrane protein (BstA), periplasmic protein (BstB), and outer membrane associated protein (BstC). Their ability to recognize and bind sterols was confirmed using protein-lipid pull down assays, where they showed selective binding to C-4 methylated sterols when in the presence of total lipid extracts from *M. capsulatus.* Quantitative assessment of ligand binding using Microscale Thermophoresis (MST) confirmed their preference for 4-monomethyl sterol, which they bind with equilibrium dissociation constants that are 30-90-fold lower than the 4,4-dimethyl sterol. High-resolution crystallographic structures of BstB and BstC reveal their pronounced divergence from eukaryotic sterol transporters. Docking studies and molecular dynamics simulations with sterol substrates reveal putative substrate binding sites and recognition mechanisms. Collectively, these data provide evidence for a novel system for sterol binding and possibly transport within the bacterial domain and advances our understanding of bacterial sterol lipids.

## Results

### Bioinformatic identification of sterol transport proteins in *M. capsulatus*

In a previous study, we used the Joint Genome Institute (JGI) Integrated Microbial Genomes (IMG) Phylogenetic Profiler to identify seventeen candidate genes in *M. capsulatus* that are unique to C-4 demethylating methanotrophs (Lee et al., 2018). Among them, a putative operon that contains five genes was found to localize next to three sterol biosynthesis genes. Two (*sdmA/sdmB*) have been characterized as demethylases involved in sterol demethylation at the C-4 position. Homologs of the other three hypothetical genes we have named *bstA, bstB* and *bstC* (<u>b</u>acterial <u>s</u>terol transporter), are found in all other aerobic methanotroph genomes that contain *sdmA/sdmB*. In some of these species, these three genes are also seen to be adjacent to other sterol synthesis genes such as the oxidosqualene cyclase (*osc*) and squalene monooxygenase (*smo*), implying they may be functionally related to sterol physiology (Fig. 1B)

To obtain additional insights into the proteins encoded by these genes, we performed more refined bioinformatic analyses including protein sequence similarity network (SSN) (Atkinson et al., 2009) and structure prediction with Phyre2 (Kelley et al., 2015) and I-TASSER (Yang et al., 2015). BstA is predicted to belong to the Resistance Nodulation Division (RND) superfamily of transporters, which comprises inner membrane proteins that facilitate drug and heavy metal efflux, and protein secretion, amongst others. They are widespread in gram negative bacteria, but homologs are also found in archaea and eukaryotes. Interestingly, the human Newman Pick Type C1, a transporter of sterols, also contains a transmembrane RND domain (Nikaido, 2018). The SSN of this superfamily shows that BstA is grouped into a single cluster containing 5 nodes (where sequences with >50% identity are merged into a single node). Structural prediction shows that BstA shares a high homology with a membrane-bound hopanoid transporter, HpnN (PDB 5KHS, *Burkholderia multivorans*), a RND transporter that moves hopanoids from the cytoplasm or cytoplasmic membrane (Doughty et al., 2011). In the SSN, HpnN is clustered separate from BstA, suggesting a possible functional divergence (Fig. 2A, Table S1).

**Figure 2.**
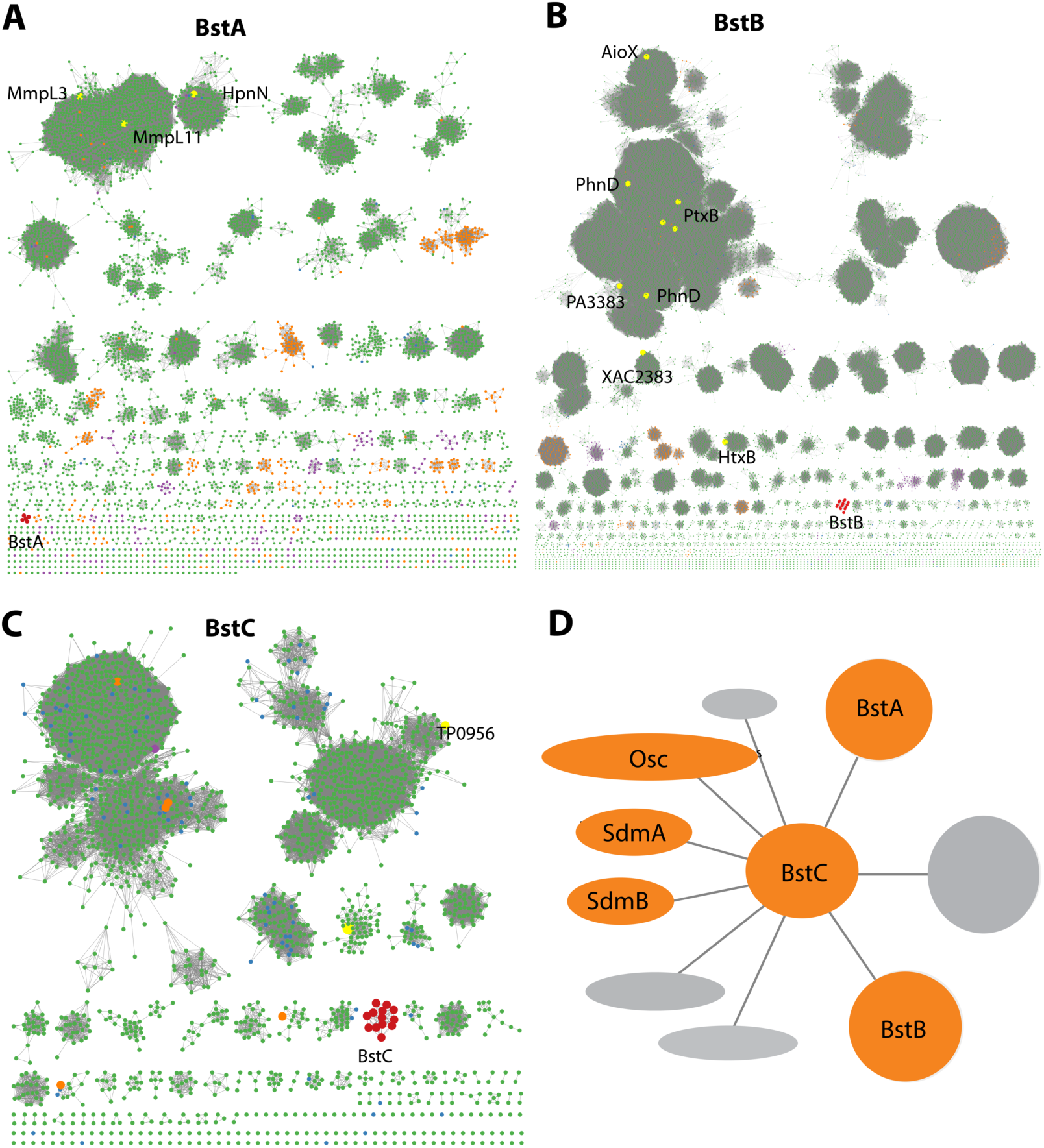
Bioinformatic analysis of Bst proteins. **(A-C)**: Protein sequence similarity networks (SSN) of BstA, BstB and BstC with the cutoff of 35%, 40%, and 40% sequence identity, respectively. Taxonomic distribution of Bst protein families showing their presence in bacterial (green), eukaryotic (purple), and archaeal (orange) genomes. **(D**) Genome neighborhood network of Bst proteins. Shown in circles and ovals are genes within 10 genes of BstC; colored in orange are those with identified or predicted roles in sterol synthesis and trafficking.

BstB belongs to a family of periplasmic/substrate binding proteins (SBPs/PBPs) known to traffic bacterial metabolites such as phosphonate. BstB proteins are grouped into a single cluster consisting of 16 sequences, with most of the sequences belonging to the *Methylococcaceae* family. Notably, the BstB proteins cluster separately even under stringent cutoffs of sequence identity (as low as 20%), hinting at a distinction of function compared with homologs (Fig. 2B, Table S1). Structure prediction of BstB shows homology to PhnD (PDB 3QK6), a periplasmic solute-binding protein of the phosphonate uptake system in *E. coli* (Alicea et al., 2011). All structurally characterized PhnD homologs are distributed into different clusters in the SSN.

Finally, BstC is predicted to be a T-component of the tripartite ATP-independent periplasmic component superfamily, which comprises periplasmic lipoproteins implicated in the cytoplasmic import of small molecules. The SSN of BstC reveals that the protein and its closest homologs are also grouped into a separate cluster even under a sequence identity cutoff of 20%. The cluster consists of 13 sequences, with most from *Methylococcaceae* and *Deltaproteobacteria* families. At the time of analysis, only one protein in the entire superfamily had structural or biochemical information: Tp0956 from *Treponema pallidum* (PDB 3U64) is found in the second largest cluster of the network (Fig. 2C, Table S1). Compared with the SSNs of BstA and BstB, that of BstC shows the family is predominantly bacterial, with only 1 and 5 sequences from eukaryotes and archaea, respectively (Fig. 2A-C).

In summary, sequence and predicted structural homology analyses reveal BstA, BstB, and BstC to be highly similar to transporters involved in disparate bacterial transport systems. However, their separation into distinct clusters in the SSNs imply their functions, including potential substrates, are not identical to their homologs. Because the Bst genes always co-localize with sterol synthesis genes (*osc, sdmA* and *sdmB*) in methanotrophs that produce sterols, we hypothesize that BstA/BstB/BstC represent a novel transport system for sterols in bacteria (Fig. 1B and Fig. 2D).

### Sterol interaction with transporters

To determine whether BstA, BstB, and BstC are sterol transporters, we first attempted gene deletion in *M. capsulatus* (data not shown). The deletion of these genes, along with others that are required for sterol synthesis (e.g., *osc*) were not successful, hinting that sterol synthesis and proper localization might be essential in this organism. Because these proteins share homology with well-known bacterial transporters, we reasoned that they are transporters and focused on determining their substrate preference by conducting pull-down assays to assess the binding of sterols from *M. capsulatus* lipid extracts. In these studies, an excess of recombinantly produced pure proteins was incubated with either the total lipid extract (TLE) from *M. capsulatus* or a specific polar fraction enriched for native hydroxy-lipids/sterols (HS). Using the HisPur Ni-NTA resin, proteins were isolated from the protein-lipid mixtures, and the protein-bound lipids were extracted and identified by GC-MS. In the presence of the TLE and HS fraction, BstA^PD^ (the periplasmic domain of BstA; details in method), BstB, and BstC bound to 4-monomethyl and 4,4-dimethyl sterols (Fig. 3A, Fig. S1). In contrast, neither the 4-monomethyl sterol, the 4,4-dimethyl sterol, nor any other lipids were present when the proteins were incubated with the DMSO negative control, suggesting that the recombinantly produced proteins did not co-purify with any lipids from the expression host. These results indicate that BstA^PD^, BstB, and BstC preferentially bind C-4 methylated sterols in a mixture of native *M. capsulatus* sterols, hopanoids, and fatty acids.

**Figure 3.**
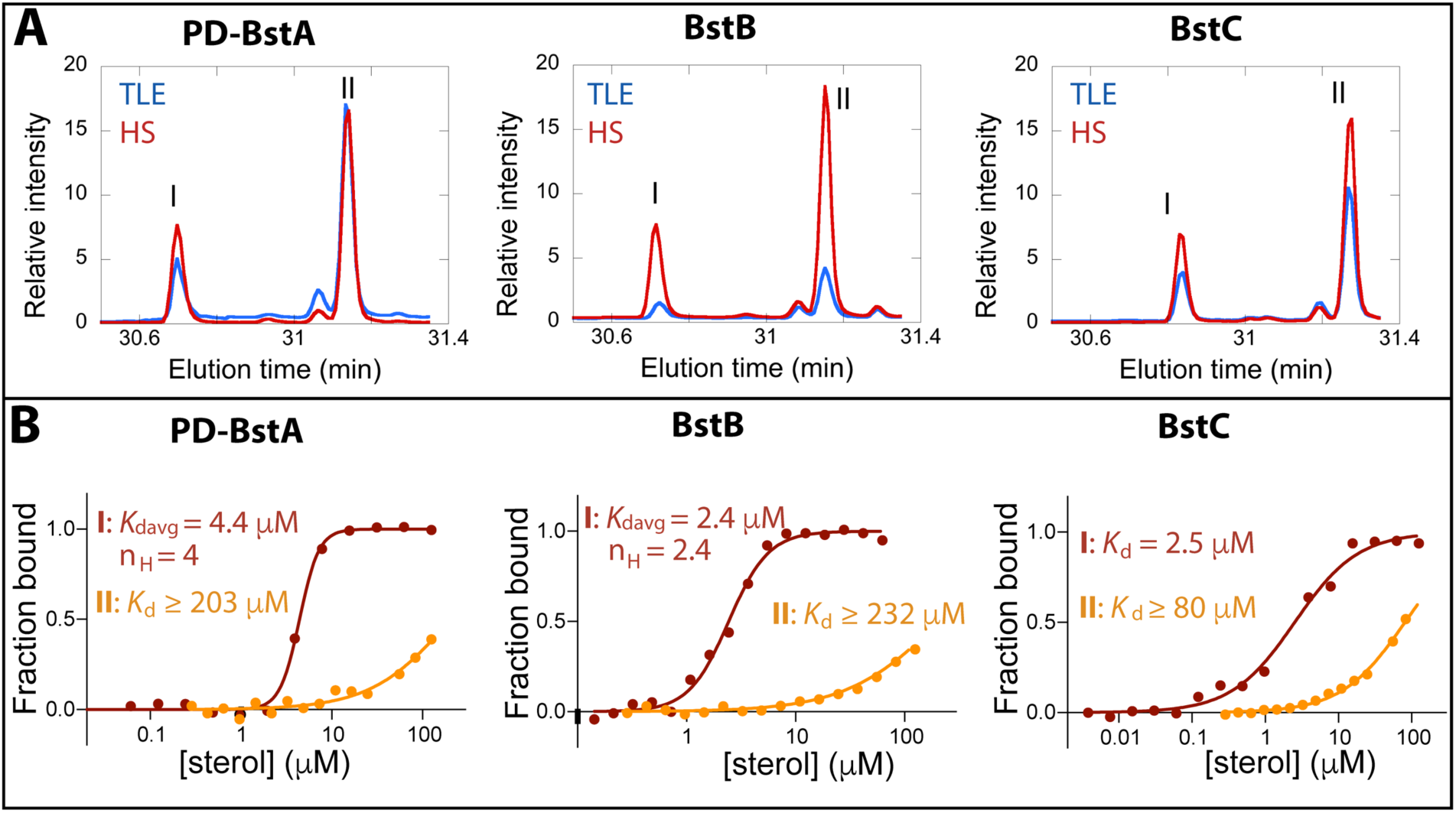
Binding of C-4 methylated sterols to transporter proteins. (A) GC chromatograms of sterols bound to purified (40 μM) *M. capsulatus* BstA^PD^, BstB, and BstC during protein-lipid pull-down assays. The proteins preferentially bind 4-monomethyl (I) and 4,4-dimethyl (II) sterols when incubated with the total lipid extract (TLE) and hydroxy lipid (HS) fraction from *M. capsulatus*. (B) MST analysis of the proteins (50 nM) binding to C-4 methylated sterols (3.8 nM -1125 mM). Equilibrium dissociation constants (*K*_d_) and Hill coefficients (n_H_) are reported on the plots.

### Quantitative analyses of protein-sterol interactions

The data from the pull-down assays demonstrated that C-4 methylated sterols have affinity for these putative sterol transporters. We next used microscale thermophoresis (MST) to determine the equilibrium binding affinities for these substrates. Binding curves were generated upon titration of the labeled proteins with serially diluted sterol substrates, and the equilibrium dissociation constants were calculated by fitting the curves with different kinetic models (Fig. 3B, methods). The “*K*_d_” model applies to binding events with a single binding site or multiple independent binding sites, while the Hill model is invoked in instances where multiple binding sites that exhibit cooperativity are present, with Hill coefficients (n_H_) greater than 1 signaling positive cooperativity. The average equilibrium dissociation (*K*_d_) constant for BstA^PD^ binding to 4-monomethyl sterol was determined to be 4.41±0.14 μM (n_H_=4.0) and ≥203±155 μM for the interaction with 4,4-dimethyl sterol. For BstB, the *K*_d_ value of 2.43±0.13 μM (n_H_ of 2.4) was determined for 4-monomethyl sterol; and ≥232±84.3 μM for 4,4-dimethyl sterol. With BstC, the determined values were 2.50±0.39 μM and ≥80.9±14.2 μM for 4-monomethyl sterol and 4,4-dimethyl sterol, respectively. All *K*_d_ values for the 4,4-dimethyl sterol could not be accurately determined due to large errors caused by non-saturation even at the highest substrate concentration. In all three proteins, we measured clear binding to C-4 methylated sterols, with marked preference for the 4-monomethyl substrate. Additionally, BstA^PD^ and BstB exhibit cooperativity for binding to 4-monomethyl substrate, indicating that multiple binding sites are plausible.

To further define the specificity of Bst proteins for sterols, we again used MST to detect their interactions with cholesterol and lanosterol, a lipid and precursor that differ in their methylation at the C-4 and C-14 positions, as well as their unsaturation patterns in the core ring structure. The MST results revealed no interaction between BstA^PD^ and BstB with either cholesterol or lanosterol. However, BstC did bind to cholesterol with a *K*_d_ value of 3.6±0.64 μM and to lanosterol with a value of 17.3±3.92 μM (Fig. S2). These data hint that BstC is more tolerant to non-native substates.

### Crystallographic structure of BstB

To understand the molecular details that govern sterol recognition and binding, we crystallized BstB and obtained crystals of the apo form (Table S2). The 1.6 Å-resolution structure of was determined by experimental phasing, producing electron density that allowed the unambiguous building of a model of BstB (except for one disordered loop spanning residues 202-204). The structure comprises two globular α/β domains (Fig. 4A, domains A and B) that form a cleft at the middle. Domain A contains six β-strands while domain B contains five. All the β-strands are flanked by α-helices. The domains are connected by two loops: one 9-residue loop that runs from β4 in domain A to β5 in domain B, and a second 8-residue loop that connects β9 in domain B and β10 in domain A. These two long loops could work as a hinge to allow a bending motion of two domains to induce a conformational change upon substrate binding that is often observed with SBPs/PBPs (Boer et al., 2019). In addition, an α-helix formed by the C-terminal residues lies behind the cleft and may also allow flexibility of the two domains. Several extend loops are found on the domain-domain interface. The loops (residues 34-38 and residues 171-177) exhibit poor electron densities, suggesting conformational sampling of more than one state (Fig. S3A).

**Figure 4.**
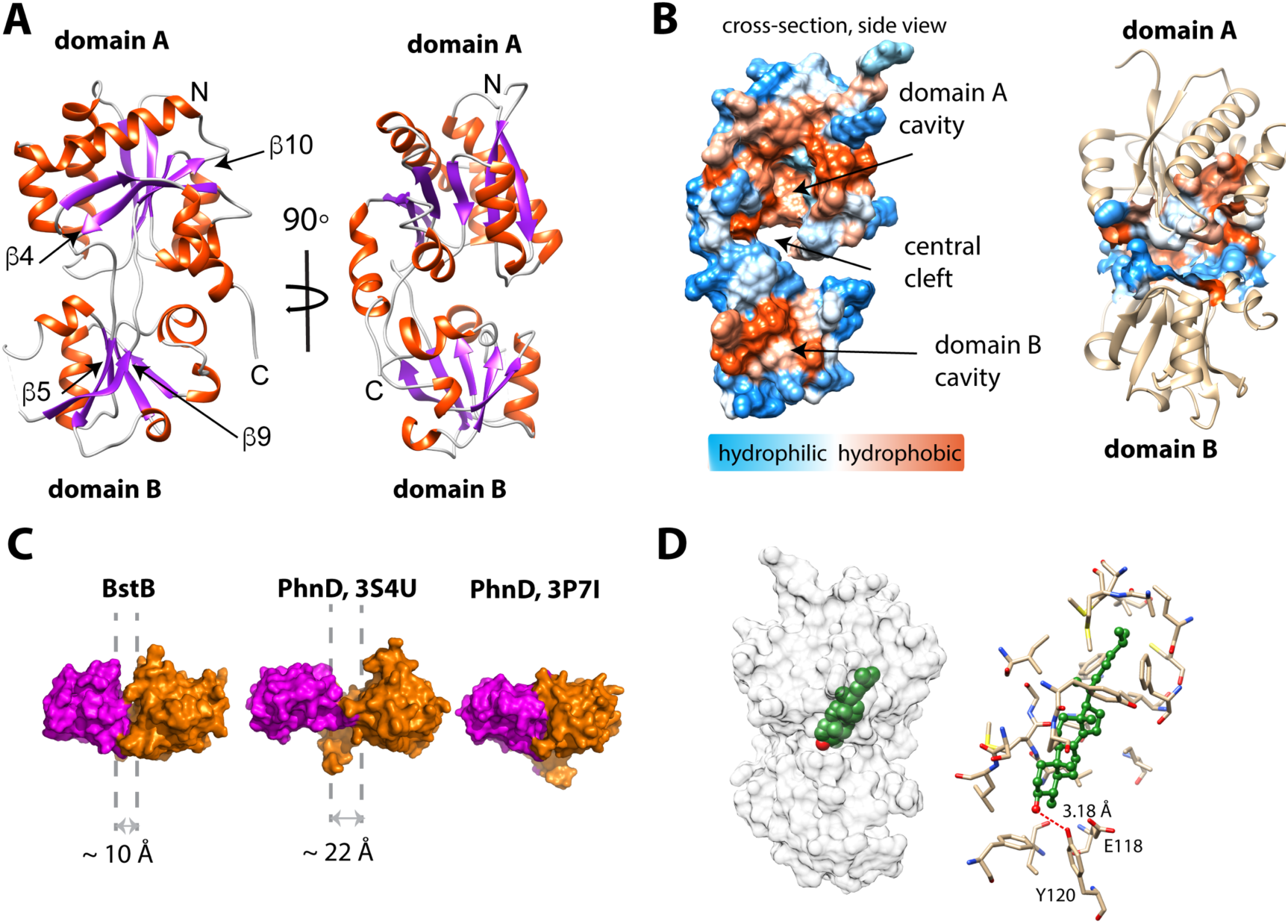
Structure of BstB. (**A**) Cartoon representation of BstB with the αhelixes and β-strands colored in red and magenta, respectively. (**B**) Hydrophobicity representations of BstB show a cleft in the middle and a cavity in domain A. The cavity is dominatingly hydrophobic and open only to one side. (**C**) Comparison of the cleft between the two domains of BstB, apo-PhnD (3S4U), and liganded PhnD (3P7I). BstB displays a similar but narrower opening (∼10 Å) compared to non-ligand PhnD (∼22 Å). (**D**) Docking model of the structures of BstB with 4-monomethyl sterol. **Left,** the docking model is shown in surface representation with 80% transparency. Sterol is shown in sphere and colored in green with the hydroxyl group is colored in red; **Right,** docking complex of *Mono_A_*. The surrounding residues are shown in stick and ball. Hydrogen bonds between the hydroxyl group of the sterol and Tyr120 is shown in the red dashed line.

The cleft between domains A and B forms a bulky cavity with an area of ∼ 1179.5 Å^2^ and a volume of ∼ 1009.6 Å^3^ that extends into domain A and is only accessible to solvent from one side of the structure. The cavity is predominantly hydrophobic, suggesting large and hydrophobic substrates like sterols are favorable for binding (Fig. 4B, Table S4). Additionally, a second pocket (area of ∼172 Å^2^; volume of ∼117 Å^3^) is found in domain B and close to the central cleft. This pocket is localized behind the unmodeled loop (residues 202-204), with the extremely poor density in this part implying high flexibility. This pocket is highly hydrophobic, but its role (if any) in ligand binding is unexplored (Fig. 4B; Fig.S4). Other separate hydrophobic areas are also observed on the protein surface and a new hydrophobic interface could be formed by two hydrophobic areas across the cleft upon conformational change induced by substrate binding (Fig. S4). These areas may be helpful for mediating the protein-protein interactions with BstA or BstC to facilitate the substrate transfer.

A DALI structural homology search (Holm and Rosenström, 2010) suggests the structure of BstB closely resembles that of PBPs that share a bi-lobe architecture and undergo conformational change upon substrate binding to the central cavity (Quiocho and Ledvina, 1996). A comparison of the BstB structure with the phosphonate-binding protein PhnD from *E. coli* with and without its substrate (2-aminoethyl phosphonate, 2AEP, PDB 3P7I and 3S4U) was performed (Alicea et al., 2011). The comparison revealed that BstB adopts an intermediate conformation between the unliganded PhnD (3S4U) and liganded one (3P7I, Fig. 4C). The cleft in BstB is less open (∼10 Å) than that in apo-PhnD (∼22 Å). There are several possible explanations for this: 1) compared with apo-PhnD structure, the cleft of BstB is deeper and more hydrophobic. Given that BstB binds to sterols (which have a nearly planar architecture and are more hydrophobic than phosphonates), the different architecture of the cleft in BstB may be related to its functional role; 2) it is known that binding of non-substrate ligands can at times trigger a conformational change in PBPs (Boer et al., 2019). Although no obvious ligand density is observed in the BstB structure, weak unmodeled densities were observed inside the central cavity. Most of these are small patches that cannot be satisfactorily modeled with water and do not closely correspond to any reagent used for crystallization. One of these patches in the central cleft could be modeled with a chloride ion engaging in polar interactions with the surrounding Glu118, Tyr120 and Phe150. Additionally, more unmodeled positive density is observed in further into the cavity in domain A (Fig. S4B-C). It is plausible that the binding of ligands co-purified with the protein triggered a conformational change leading to a “partially-open” state. A third possibility is that BstB is structurally distinct from the PhnD homologs in that its cleft is much smaller.

### Substrate docking and molecular dynamics simulation of BstB

To obtain greater insights into the mechanism of substrate recognition and binding in the absence of a substrate-bound structure, we used ICM-Pro (Abagyan et al., 1994; Abagyan and Totrov, 1994) to generate docking models of C-4 methylated sterol binding to BstB. A three-dimensional model of 4α-monomethyl sterol was constructed from the structure of cholesterol using the Molecular Editor in ICM-Pro and then docked to the BstB structure. In the first iteration, the protein was held rigid, and the ligand allowed to dock flexibly. In this docking model, the sterol was positioned in the hydrophobic cavity in domain A with the polar head pointing toward the central cleft and domain B, adjacent to three residues: Glu118, Tyr120 and Asn192. In subsequent replicate docking iterations, the side chains of these three residues were allowed to be flexible along with the ligand. There are three energetically favored poses: two (*Mono*_A_ and *Mono*_B_) form a single hydrogen bond to Tyr120, while pose *Mono*_C_ where the hydroxyl makes two hydrogen bonds to Tyr120 and Asn192 (Fig. 4D and Figs. S5A-C, Table S3).

To probe the dynamics of the BstB structure during substrate binding, we performed multiple 20 ns molecular dynamics (MD) simulations using Desmond (Schrodinger) on each docked poses for monomethyl sterol from ICM-Pro. In all cases, analysis of the resultant MD trajectories indicated that the simulations became stable after the first few picoseconds. In the three poses where there was initially an interaction with Tyr120, that interaction is lost and replaced with a new hydrogen bond with Glu118, which persists for 85%-95% of the simulation. Pose *Mono*_C_ also has an intermittent interaction with Asn192 for approximately 11% of the simulation. In all simulations, whether there is a hydrogen bonding interaction with the ligand or not, the amide group of Asn192 is flipped.

To better probe the protein’s substrate preference, docking and MD simulations with the 4,4-dimethy sterol were also generated. In these models, the 4,4-dimethyl sterol also docks to the cavity in domain A with three poses of roughly equal energies (Figs. S5D-F, Table S3). In one (pose *Di*_A_), hydrogen bonds to Glu118, Tyr120 and Asn192 are possible; in a second (*Di*_B_), hydrogen bonds to Glu118 and Tyr120 are possible; the third (*Di*_C_) has a hydrogen bond to Tyr120 only. The 4,4-dimethyl sterol showed interesting behavior during the 20 ns simulations. Pose *Di*_A_, which had three hydrogen bonds after ICM-Pro docking, retained all three interactions (although those with Glu118 and Tyr120 are intermittent and persist for only 12% of the simulation). Pose *Di*_B_ retained both interactions from the docking model, although the hydrogen bond to Tyr120 was only present for 10% of the simulation. The third pose, *Di*_C_, showed the largest structural changes, losing the one interaction it had with Tyr120. In this pose, the head group of the sterol moved out of the binding pocket delineated by Glu118, Tyr120 and Asn192 and a new hydrogen bond with the carbonyl oxygen of Pro34 formed, persisting for the duration of the simulation (Table S3). It is possible that these differences, although subtle, explain the reduction in affinity for the dimethyl sterol substrate.

The aliphatic tails of all docked sterols reside in a hydrophobic cavity in domain A lined by the side chains of Tyr33, Pro34, Ala46, Met47, Met50, Ile89, Ser91, Leu92, Pro110, Ile112, Leu219, Met220, Val222, Leu244, Leu253, Cys254, Phe257, Ile259 and Phe262 in domain A. Sequence alignment shows that these residues are highly conserved between proteins in the same SSN cluster, implying their significance in forming the substrate binding pocket. Mutagenesis to remove or reduce the potential interactions with the polar head and the aliphatic tail showed subtle influences (<2-fold) on the sterol binding affinity, indicating that the interaction between the protein and sterol is extensive and principally depends on hydrophobic interactions that are not easily disrupted (Fig. S7). Moreover, the conformational dynamics of PBPs could mask the identity of additional residues that might be involved substrate binding and stabilization. Regardless, the determination of the sterol binding site in BstB remains to be experimentally verified.

To investigate whether BstB can undergo conformational change (as most PBPs do upon substrate binding), a 150 ns molecular dynamics simulation on apo-BstB was performed. Plotting the *RMSD* of the simulated structure against the initial crystal structure at 25 ps intervals during the MD trajectory shows that for the first ∼90 ns the protein showed some degree of flexibility (Fig. S8A). During this part of the trajectory, the *RMSD* oscillates between 2.0 to 3.1 Å (average *RMSD* =2.6 Å). At ∼90 ns, there was a distinct conformational change in the structure giving rise to conformations with *RMSD*s ranging from 2.7 to 4.5 (average *RMSD* =3.3 Å). Visual analysis of the MD trajectory shows that at ∼90 ns the two domains open by approximately 20° (Fig. S8B, Movie 1). In contrast, the simulation (150 ns) on the *Mono*_C_ docked complex which had a *RMSD* of 2.6 Å for the entire length of the trajectory and did not show the same conformational change as the apo-BstB simulation, which supports the idea that the apo-BstB structure represents a “partially-open” form of the protein (Movie 2).

### Crystallographic structure of BstC

The structure of BstC was also determined by experimental phasing to a resolution of 1.9 Å. Two monomers (*RMSD* of 0.656 Å for all Cαs) in the asymmetric unit form a dimer with 180^∘^ rotational symmetry. Each monomer adopts an all-α-helical structure with 12 α-helices in total, designated with H1 to H12; 9 antiparallel arranged helices (H4-H12) form an irregular ring with a pore running through the center. The H4, H6, H8, H10, H12 constitute the inside wall of the pore, while the others form the outside wall. Two helices (H1-H2) at the N-terminus sit on the side-top of the ring with a short helical turn (H3) connecting the two (Fig. 5A).

**Figure 5.**
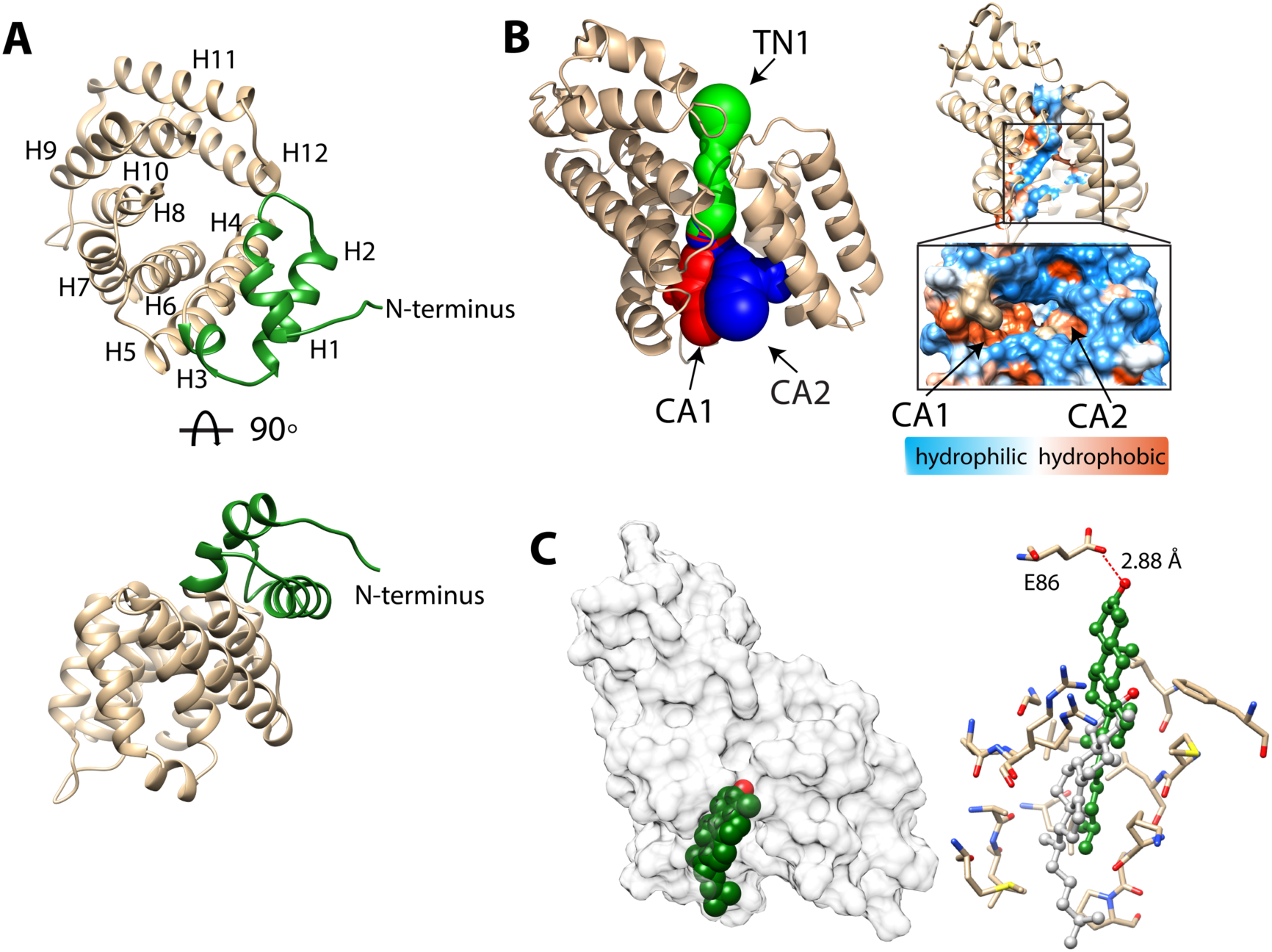
Structure of BstC. (**A**) Cartoon representation of BstC. The two αhelixes at the N-terminus are colored in green. Two monomers (RMSD of 0.656 A for all Cαs) with 180° rotational symmetry exist in the asymmetric unit. (**B**) A representation of cavities BstC (left, with Tunnel 1 colored in green, Cavity 1 in red, and Cavity 2 in blue) alongside hydrophobicity representation of the cavities (right). (**C**) Docking model of the structures of BstC with 4-monomethyl sterol. **Left,** the docking model is shown in surface representation with 80% transparency. Sterol is shown in sphere and colored in green with the hydroxyl group is colored in red; **Right,** docking complex before and after MD simulation. The surrounding residues are shown in stick and ball. The sterol prior to the MD simulation is colored in grey and while green shows its position after the simulation. A hydrogen bond between the hydroxyl group of the sterol and Glu86 is shown as a red dashed line.

Hydrophobicity analysis shows that BstC has a predominantly hydrophilic exterior surface (Fig. S9A). Three cavities are found in the center of the structure. We annotated them as Tunnel 1 (TN1), Cavity 1 (CA1) and Cavity 2 (CA2), with total area of ∼ 597.1 Å^2^ and total volume of ∼ 447 Å^3^. TN1 is closer to the N-terminal face of the structure, while CA1 and CA2 are proximal to the C-terminal face and form two open hydrophobic pockets; TN1 exhibits a mixture of hydrophobic and hydrophilic amino acids (Fig. 5B and Fig. S8B, Table S4). The extensive hydrophobic environment inside the BstC structure suggests it can accommodate large hydrophobic substrates. Notably, electrostatic calculation shows that most of the surface of BstC, especially the C-terminal face, exhibits strong negative charge (Fig. S9). These charged surfaces may mediate protein-protein interactions, perhaps with BstB.

A structure similarity search using the DALI server revealed high homology to Tp0956 (PDB 3U64, *Treponema pallidum strain Nichols*). Tp0956 is the T-component protein of a subfamily of TRAP (tripartite ATP-independent periplasmic) transporters known as TPATs (tetratricopeptide repeat-protein associated TRAP transporters), which are implicated in the transport of organic acid ligands in bacteria (Brautigam et al., 2012). The true physiological substrate of Tp0956 has not been determined, but *Treponema* is a spirochete known to rely on import of host lipids and fatty acids to construct its cell envelope (Radolf et al., 2016). Tp0956 contains four helical hairpins that are similar to tetratricopeptide repeat (TPR) motifs, which are hallmarks of proteins that are involved in protein-protein interactions (D’Andrea and Regan, 2003). Comparison of the structures show that two antiparallel α-helices exist in Tp0956 are missing in the structure of BstC (between H5-H6), as well as a shorter H4 and H5. But the TPR-like substructure is also observed in BstC and forms part of the pore and lateral surface of the protein, hinting that BstC might engage with itself or other proteins (Fig. S10). Size-exclusion chromatography coupled with multiangle light scattering (SEC-MALS) analysis shows that indeed the purified BstC exists both as a monomer and a dimer in solution with a ratio ∼ 3:1 (Fig. S11). However, the role of the dimerization in this protein is not clear and its determination will require additional studies.

### Substrate docking and molecular dynamics simulations with BstC

To assess substrate-binding, we performed the docking of 4-monomethyl sterol and 4,4-dimethyl sterol into the BstC structure using ICM-Pro. The final docking model was obtained by choosing the one with the most negative ICM-Pro score. All the sterols were found embedded in the hydrophobic cavity (CA1) lined by Gly128, Val131, Arg132, Met135, Ala138, Leu139, Leu142, Leu145, Leu167, Met170, Leu171, Lys174, Ala175, Pro176 and Phe208, with the hydroxyl group pointing inwards and the aliphatic tail pointed out (Fig. 5C and Fig. S12A-B). Residues involved in sterol interaction are located on H6, H7, H8 and H10 and are all highly conserved in a sequence alignment of the closest BstC homologs. No hydrogen bond is anticipated between the sterol and protein in the docking models.

We then performed multiple 20 ns molecular dynamics simulations on the docked models of 4-monomethyl sterol from ICM-Pro. The simulations reveal that the substrate moves approximately 8-9 Å further into the binding site and the polar head makes a hydrogen bonding contact with the side chain of Glu86 (Fig. 5C). This movement takes place almost immediately and the hydrogen bond persists for 94% of the total time. In both the docking models and MD simulations, the sterol is positioned near the entrance of CA1 from the C-terminal side. Next to the entrance is an extended loop that consists of highly conserved hydrophobic and neutral residues (residues 175-184).

Intriguingly, simulations with the 4,4-dimethyl sterol revealed that it behaves differently in that it moves slowly inwards and does not get as deep (∼2.6 Å) into the pocket as the monomethyl sterol (Fig. S12C). A water-mediated contact is formed between the polar head and the side chain of Glu86 but persist for only 2% of the time. This lack of additional movement and H-bonding interaction may explain the binding data, where the protein has a *K*_d_ for the monomethyl sterol that is ∼ 30× lower than for the dimethyl sterol substrate.

Mutagenesis to convert Glu86 to Gly and Trp to remove or reduce the potential interactions with the polar head of the sterols leads to ∼ 4.5- and 100-fold decrease in the equilibrium affinity for 4-monomethyl sterol (Fig. S143A). Meanwhile, the mutagenesis eliminates the binding to 4,4-dimethyl sterol, cholesterol and lanosterol (Fig. S13B,C). These findings indicate that Glu86 plays a critical role in interacting with and stabilizing the sterols.

### Comparison to eukaryotic sterol transporters

In most eukaryotic cells, lipid transfer proteins (LTPs) are used to transfer sterols between subcellular membranes by non-vesicular transport to maintain sterol homeostasis in different sub-cellular organelles.(Baumann et al., 2005) The most-studied family in mammals is the steroidogenic acute regulatory protein (StAR)-related lipid transfer (START) domain (STARD) proteins. All fifteen STARD proteins identified to date possess lipid-harboring START domains that can be divided into six subfamilies based on sequence similarity and ligand specificity. In addition to STARD proteins, a novel yet evolutionarily conserved family that contain the START-like domains called the GRAMD1s/Lam/Ltc family (GRAMD1s in mammals and Lam/Ltc in yeast) was shown to mediate non-vesicular sterol transport from the plasma membrane to the endoplasmic reticulum to maintain sterol homeostasis (Gatta et al., 2015). A third family consists of the Osh/ORP/OSBP proteins (Oxysterol-binding homology in yeast and Oxysterol-binding Protein/OSBP-Related Protein in mammals), which are considered to be either sterol-sensing and/or sterol-transfer proteins (Raychaudhuri and Prinz, 2010). Besides these, two Niemann-Pick type C proteins, NPC1 and NPC2, were also reported to directly bind and transport cholesterols and various oxysterols; defective NPC proteins causes NPC disease, a fetal neurodegenerative disorder where sterols aberrantly accumulate in the lysosome (Carstea et al., 1997).

To better understand the differences between eukaryote and bacteria sterol transport, we compared the structures of BstB and BstC with well-studied eukaryotic sterol transporters from different families. Human STARD4 (PDB 6L1D), yeast Lam4 (PDB 6BYM), yeast Osh4 (PDB 1ZI7 and 1ZHY), yeast NPC2 (PDB 6R4M and 6R4N), as well as a non-specific cytosolic sterol carrier protein, rabbit SCP2 (PDB 1C44) were chosen for the comparison. The crystal structures of STARD4, Lam4 and Osh4 show that they share a similar fold in that they are dominated by β-sheet motifs, which wrap around the longest α-helix and fold into a half-barrel/groove structure with the other side (or two) packed with several α-helices (Fig. S14A-D). A central hydrophobic cavity can accommodate sterols. In most cases, the sterols are positioned proximal to the tunnel entrance with the hydroxyl end oriented towards the inside of the tunnel. Also, a “lid” comprising a flexible “Ω1” loop is observed near the entrance of tunnel in STARD4 and sterol-bound Lam4. This loop is replaced by an N-terminal α-helix in Osh4, which serves a similar function. The SCP2 structure also exhibits an α/β-fold with a “lid”-like α-helix observed close to the N-terminus (Fig. S14E). The sterol-bond NPC2 adopts a different immunoglobulin-like β-sandwich fold with seven strands arranged into two β-sheets with a loosely packed hydrophobic cavity formed between the two sheets. The sterol is positioned at the entrance of the cavity with the hydroxyl end pointing out; no “lid”-like structure is observed in NPC2 structure (Fig. S14F-G).

Our comparative analysis shows that the eukaryotic sterol transporters share very little resemblance with BstB and BstC. The bilobed architecture of BstB is pronounced in its divergence from eukaryotic sterol transporters, suggesting that this protein evolved separately and may work in a distinct manner. For the all-helical BstC, the only similarities are the presence of a hydrophobic tunnel formed in the center of the protein that can accommodate sterols, and a loop next to the tunnel’s entrance that may also be used as a “lid” to restrict access to the substrate binding site (Fig. S14H).

## Discussion

Since their discovery more than 40 years ago, the function of bacterial sterols has remained a mystery. Recent identification of bacterial sterol demethylase enzymes that diverge from their eukaryotic counterparts provided renewed enthusiasm for deciphering the molecular machinery involved in bacterial sterol synthesis and trafficking (Lee et al., 2018). The aerobic methanotroph *M. capsulatus* is known to synthesize 4-monomethyl sterol from 4,4-dimethyl sterol and to accumulate these sterols in its outer membrane. Until now, the transport process remained elusive owing to lack of knowledge of transporters that facilitate the trafficking. We now fill this gap with the identification of potential transporters of C-4 methyl sterols in *M. capsulatus*: BstA, BstB and BstC.

Bioinformatic analyses revealed that BstA, BstB, and BstC belong to well-studied families of bacterial metabolite transporters. High resolution structures of BstB and BstC show that their large ligand binding sites display a dichotomy of hydrophobic and hydrophilic amino acids whose sidechains interact with the sterol hydrophobic tail and hydrophilic head. For BstC, this is especially interesting, as the only other family member characterized, Tp0956 from *Treponema pallidum*, displays a similar dichotomous ligand site. *Treponema pallidum*, the causative agent of syphilis, is a spirochete known to acquire and use host lipids– it is possible that Tp0956, and perhaps other T-component proteins, are involved in lipid trafficking.

In summary, the Bst proteins share sequence and structural homology with bacterial transporters involved in the trafficking of essential nutrients such as sugars, amino acids, vitamins, solutes, and metal ions (Davidson et al., 2008; Higgins, 1992). Their implication in the transport of sterols expands the substrate repertoire for their respective families. The substrate binding sites detected in BstB and BstC should now enable the curation of functionally homologous sites in bacterial proteins, including in those from human pathogens known to traffic host lipids.

## Materials and methods

Detailed materials and methods describing bioinformatic analyses are provided in an extended materials and methods section of the Supporting Information. Also provided are methods for recombinant protein production and purification, lipid extraction form *M. capsulatus*, protein-lipid pull-down assays and chromatographic analyses, MST experiments, macromolecular crystallography, molecular docking, and simulations.

## Acknowledgments

This work was supported in part by NSF grant MCB1919153 (to L.M.K.D. and P.V.W.). L.M.K.D. was supported by funding from the Terman, Gabilan, and Hellman Fellowships, and is a MAC3 Impact Philanthropies Faculty Fellow. Crystallographic data were acquired at the Stanford Synchrotron Radiation Lightsource. Use of the Stanford Synchrotron Radiation Lightsource, SLAC National Accelerator Laboratory, is supported by the U.S. Department of Energy, Office of Science, Office of Basic Energy Sciences under Contract No. DE-AC02-76SF00515. The SSRL Structural Molecular Biology Program is supported by the DOE Office of Biological and Environmental Research, and by the National Institutes of Health, National Institute of General Medical Sciences (P30GM133894). The contents of this publication are solely the responsibility of the authors and do not necessarily represent the official views of NIGMS or NIH.

