## Supporting Information for "Novel sterol binding domains in bacteria"

### Materials and Methods

#### Bioinformatic analyses

The SSNs were generated using EFI-EST sever (http://efi.igb.illinois.edu/efi-est/stepa.php). Proteins are represented by nodes. Only the nodes that share pairwise sequence alignments greater than the user-specified value will be connected by edges to create a network that reveals potential functional relationships among the proteins (Atkinson et al., 2009). HMMER webserver (https://www.ebi.ac.uk/Tools/hmmer/) was used to predict the superfamilies of Bst proteins. BstA is predicted to be a member of MMPL family (pfam03176 or IPR004869). Because some homologs are not annotated in the pfam family, we first used the Blast function to identify proteins sharing homology with BstA in UniProtKB (www.uniprot.org). Homologs within the cutoff value of $5.3 \times 10^{-80}$ were also used in the SSN creation. The SSN was generated using the EFI-EST online tool (Gerlt et al., 2015) with an E-value threshold of $10^{-60}$ (~35% sequence identity); with this threshold, BstA proteins were grouped into a single cluster. UniRef50 (>50% sequence identity merged into a single node) was used to facilitate the calculation and save time during the SSN creation. BstB belongs to a family of ABC transporter phosphonate periplasmic substrate-binding proteins (pfam12974); this Pfam was selected to create an SSN at E-value threshold of $10^{-50}$ (~40% sequence identity). Homologous proteins found in the UniprotKB (cutoff at $1.6 \times 10^{-62}$) were added to the network as well. UniRef90 standard (≥90% sequence identity merged into a single node) was used to create the BstB SSN. BstC is classified as a member of the TRAP transporter T-component superfamily (IPR038537), which are thought to facilitate the import of small molecules into cytoplasm (Radolf et al., 2016). An SSN was constructed with this dataset and the homologous genes found in the UniprotKB (cut-off at $6.1 \times 10^{-39}$) at E-value threshold of $10^{-55}$ (~40% sequence identity). Cystoscope software (Shannon et al., 2003) was used to visualize the SSNs with the “organic” layout. The genome neighborhood network and diagram were generated using the Enzyme Function Initiative-Genome Neighborhood Tool (EFI-GNT; https://efi.igb.illinois.edu/efi-gnt/) with the SSNs generated above. The neighborhood reading frame was set to 10 frames and the minimal co-occurrence was set to 20%.

### Attempted deletion of *M. capsulatus* Texas H156DRAFT_2759-2757

In attempts to generate the *M. capsulatus* Texas H156DRAFT_2759-2757 mutant, a linear DNA fragment was PCR-amplified from a plasmid containing a kanamycin resistance gene flanked by 1kb *M. capsulatus* chromosomal sequence upstream of H156DRAFT_2759 and 1kb downstream of H156DRAFT_2757. Electrocompetent *M. capsulatus* cells were prepared via the following protocol: a 300 mL culture was grown to mid-exponential phase, washed three times in 5% cold sterile glycerol and resuspended in 300μL cold sterile 10% glycerol. The electrocompetent cells were mixed with 2.5 μL of the linear DNA fragment (1000 ng total) and shocked using various electroporation settings (Bacteria 1,2,3, and 5) on the Bio-Rad Pulser Xcell and the Ec1 setting on the Bio-Rad Micropulser [parameters were a combination of different cuvette gaps (0.1cm or 0.2cm), voltages (1.8-3.0 kV), resistances (200-400 Ω)]. Electroporated cells were grown up for 24hrs in 500 μL of media made with nitrate mineral salts (NMS) and methane; following this, the cells were plated on NMS-agar plates supplemented with kanamycin. No colonies were ever observed, suggesting that depletion of these transporters is toxic to the cells.

### Protein expression and purification

The genomic DNA of *M. capsulatus* Texas was subjected to several rounds of PCR to isolate the genes encoding BstB and BstC without their respective signal peptides (cleavage sites determined by the SignalP 5.0 server (Armenteros et al., 2019)). The final gene products were assembled into pET-28a vectors containing TEV-cleavable N-terminal hexa-histidine (His-) tags or pET-20b vectors containing C-terminal His-tags. DNA sequencing of the plasmids confirmed the correct insertion of the genes. The $BstA^{PD}$ gene was synthesized and inserted into pET-28a after codon optimization for expression in *E. coli*. The soluble periplasmic domain of BstA ($BstA^{PD}$) was designed based on the structure predicted by AlphaFold (Jumper et al., 2021). Briefly, two fragments spanning residues 54-277 and 510-732 were connected by a GSGSGSGS linker to replace the membrane-spanning sequence. The DNA encoding $BstA^{PD}$ was cloned into pET28a. All the protein sequences and primers are listed in Table S5.

BstA$^{PD}$, BstB (residues 24-281) and BstC (residues 16-264) proteins were expressed in *E. coli* BL21(DE3) cells and grown at 37 °C until $OD_{600nm}$ of 0.6. Protein expression was induced upon the addition of 0.25 mM isopropyl β-D-1thiogalactopyranoside (IPTG) while shaking at 18 °C for 12h. Cells were harvested through centrifugation at 8000 × g for 15 min and resuspended in lysis buffer containing 25 mM Tris-HCl (pH 8.0), 100 mM NaCl, 2 mM 2-mercaptoethanol, and 1 mM phenylmethylsulfonyl fluoride (PMSF). Cells were lysed by microfluidization and centrifuged at 11,000 × *g* for 40 min to remove cell debris. The supernatant was applied to a Ni-NTA affinity column (HiTrap IMAC FF, Cytiva product # 17092104), washed with lysis buffer supplemented with 15 mM imidazole to remove weakly bound proteins, and eluted with lysis buffer containing 300 mM imidazole. The eluate was concentrated and applied to a size-exclusion column (HiLoad 16/600 Superdex 200 pg, Cytiva product # 28989335) to remove additional contaminants, and purity was assessed through gel electrophoresis.

Selenium-methionine (Se-Met) labeled BstB and BstC were produced by supplementing the endogenous methionine synthesis of the expression host with Se-Met. The cells were cultured in M9 minimal media until $OD_{600nm}$ of 0.4. L-lysine (100 mg/L), L-phenylalanine (100 mg/L), L-threonine (100 mg/L), L-isoleucine (50 mg/L), L-leucine (50 mg/L) and L-valine (50 mg/L) and L-Se-Met (60 mg/L) were added to the media. 0.25 mM IPTG was added after 1h to commence induction and cells were cultured for an additional 12h at 18°C before harvesting. The proteins were purified by the protocol described above.

The N-terminal His-tags of proteins were removed during overnight incubation at room temperature with TEV protease at a molar ratio of 1:100 (TEV: protein). Proteins without tags were further purified by size-exclusion chromatography. The molecular weight of proteins was determined by SEC-MALS (Wyatt Technology). The purity of all purified proteins was assessed using SDS-PAGE stained with Coomassie Brilliant Blue, after which aliquots of the purest fractions were flash-cooled in liquid nitrogen and stored at -80 °C.

**Size-exclusion chromatography (SEC)-coupled multi-angle light scattering (MALS)**

SEC-MALS was performed using an in-line Superdex 200 Increase 3.2/300 GL SEC column (GE Healthcare) combined with a miniDawn Multi-Angle Light Scattering (MALS) detector coupled with an Optilab refractive index detector (Wyatt Technology, Santa Barbara, CA, USA). 15 μL (~ 200 ng) protein was centrifuged at 13000 × g for 10 min before injected into the pre-equilibrated SEC column with buffer 25 mM Tris-HCl (pH 8.0), 100 mM NaCl and 2 mM 2-mercaptoethanol. Proteins were separated at a flow rate of 0.15 mL/min at room temperature. Molecular masses were calculated using the Astra6.1 software (Wyatt Technology).

**Sterols extraction from *M. capsulatus***

Lipids were extracted from cell pellets of 2 L *M. capsulatus* cultures using a modified Bligh-Dyer extraction method as previously described.(Lee et al., 2018) Cells were re-suspended in 10:5:4 (vol:vol:vol) methanol: dichloromethane (DCM):water and sonicated for 1 hour. The organic phase was separated with 1:1 (vol:vol) DCM:water, later washed three times with DCM and then transferred to a clean vial and dried with $N_2$ to yield the total lipids extraction (TLE). The TLE was further purified using Silica column chromatography to isolate the alcohol soluble lipids.(Summons et al., 2013) The TLE was added to a hand-packed silica column and lipids were eluted using solvent solutions of different polarities (Hexane, 8:2 Hexane:DCM, DCM, 1:1 DCM:Ethyl Acetate, Ethyl acetate). The alcohol soluble lipids (HS fraction) were eluted with 1:1 DCM:Ethyl Acetate. The HS fraction was then dried with $N_2$ and dissolved in DMSO. The 4-methylsterol and 4,4-dimethylsterol were purified from the alcohol soluble lipids by liquid-chromatography (LC) using an Agilent 1260 Infinity II LC System. Briefly, lipids were run through a InfinityLab Poroshell120 EC-C18 column (4.6 x 150 mm, 2.7 Micron). The solvent system consisted of Solvent A: MeOH (with .04% formic acid and 0.1% ammonia), Solvent B: acetonitrile, and Solvent C: Water (with .04% formic acid and 0.1% ammonia). The solvent gradient started with 84% Solvent A, 1% Solvent B, and 15% Solvent C and ended with 100% Solvent A over the course of 50 minutes. Isolated sterols were derivatized to trimethylsilyl ethers with 1:1 (vol:vol) Bis(trimethylsilyl)trifluoroacetamide: pyridine and confirmed using GC-MS analysis as described in Lee et al, PNAS 2018.

**Protein-lipid pull down assay**

Recombinantly expressed and purified proteins (BstA$^{PD}$, BstB, BstC) were incubated with two different *M. capsulatus* lipid extracts: total lipid extractions (TLE) or the enriched sterol fraction of the TLE (HS), both dissolved in DMSO. The TLE (139.5 µg total) contained 65.7 ng of 4-monomethyl sterol and 338.9 ng of 4,4-dimethyl sterol. The HS fraction (99.5 µg total) contained 313.8 ng and 967 ng of 4-monomethyl sterol and 4,4-dimethyl sterol, respectively. The experimental reaction conditions are as follows: 40 µM protein (BstA$^{PD}$, BstB, BstC) was incubated with 7.5 µL of either the TLE or HS lipids. The reactions were carried out in reaction buffer of 200 mM HEPES pH 8.0, 100mM NaCl and 0.05% Triton X-100. During the negative control reactions, the proteins were incubated with DMSO that did not contain any lipids. This serves as (1) a vehicle control since the *M. capsulatus* TLE and HS fractions were dissolved in DMSO and (2) to determine whether C4-methylsterols were bound to the Bst proteins as a result of protein expression in *E. coli*. The reactions (both control and experiment) were incubated at 25 °C for 20 hours. The proteins were purified from the reactions using HisPur Ni-NTA his resin (Thermos Fisher), lipids from the eluted protein fractions were extracted using a modified Bligh Dyer protocol (Ekiert et al., 2017; Lee et al., 2018) and lipid samples were analyzed by GC-MS as previously described. HisPur Ni-NTA wash buffer: 25 mM Tris pH 8.0,100 mM NaCl and elution buffer: 25 mM Tris pH 8.0,100 mM NaCl, 300mM imidazole. SDS PAGE electrophoresis was conducted to demonstrate that the fractions analyzed did in fact contain protein.

**GC-MS analysis of *M. capsulatus* lipids**

Derivatized lipids were separated with an Agilent 7890B Series GC equipped with 2 Agilent DB-17HT columns in tandem (each 30 m x 0.25 mm i.d. x 0.15 µm film thickness) with helium as the carrier gas at a constant flow of 1.1 ml/min. The program was as follows: 100°C for 2 min, then 12°C/min to 250°C and held for 10 min, then 10°C/min to 330°C and held for 17.5 min. 2 µL of each sample was injected in splitless mode at 250°C. The GC was coupled to an Agilent 5977A Series MSD with the ion source at 230°C and operated at 70 eV in EI mode scanning from 50 to 850 Da in 0.5 s. Lipids were identified based on their retention time and compared to previously published spectra (Wei et al., 2016).

**Microscale thermophoresis**

Microscale thermophoresis (MST) experiments were performed with NanoTemper Monolith NT.115 instrument according to the manufacturer's instructions (NanoTemper Technologies). In brief, His-tagged BstA$^{PD}$, BstB and BstC were fluorescently labeled with Monolith His-Tag Labeling Kit RED-tris-NTA 2$^{nd}$ Generation according to the manufacturer's instructions. The 4-methylsterol and 4,4-dimethylsterol were extracted from *M. capsulatus*, while lanosterol (CAS:79630, Sigma-Aldrich) and cholesterol (CAS: AAA1147018, Fisher Scientific) were prepared in DMSO with a final stock concentration of 3.9 mM and 7.76 mM, respectively.

50 nM each of labeled proteins was diluted in assay buffer (20 mM HEPES pH 8.0, 100 mM NaCl, and 0.05% Tween 20), in the presence of sixteen-step serial dilution of 4-methylsterols or 4,4-dimethylsterols (ligand solubilized in DMSO at a final constant concentration of 5%). After incubation, samples were loaded into Monolith NT.115 Premium Capillaries (NanoTemper Technologies) and measurements were taken at a constant temperature of 23 °C. MST traces were collected with an LED excitation power of 40% and an MST laser power of 40%. The MO. Control Analysis software (NanoTemper) was used to analyze the interaction affinity and the equilibrium dissociation constant ($K_d$) for each ligand using the $K_d$ fit model or Hill fit model. The data was exported and plotted in Prism.

Equation of $K_d$ -model fitting:

$$f(c) = \frac{c + c_T + K_d - \sqrt{(c + c_T + K_d)^2 + 4\,c\,c_T}}{2\,c_T}$$

f(c) is the fraction bound at a given ligand concentration c. Kd is the dissociation constant or binding affinity and $c_T$ is the final concentration of target in the assay.

Equation for Hill-model fits:

$$f(c) = Unbound + \frac{Bound - Unbound}{1 + \left(\frac{EC_{50}}{c}\right)^{n_{Hill}}}$$

f(c) is the fraction bound at a given Ligand concentration c. Unbound is the $F_{norm}$ signal of the Target. Bound is the $F_{norm}$ signal of the Complex. $EC_{50}$ is the half-maximal effective concentration and $n_{Hill}$ is the Hill coefficient. The Hill coefficient $n_{Hill}$ describes the degree of cooperativity of an interaction: $n_{Hill}>1$ indicates positive cooperativity, while $n_{Hill}<1$ indicates negative cooperativity.

**Crystallization of BstB and BstC**

Sparse-matrix crystallization screens were performed with purified proteins at concentrations of about 5 mg/mL for each. All crystals were obtained by the sitting-drop vapor diffusion method by mixing 0.5 μL protein and 0.5 μL precipitant solution at 20°C. Native and Se-met BstB crystals appeared in the MCSG1-C12 condition containing 25% (v/v) PEG3500 and 0.1 M Bis-tris pH 6.5 after 1 week of incubation at 20 °C. Crystals were cryoprotected in the same conditions with the addition of 20% (v/v) PEG400 before being flash-cooled and stored in liquid nitrogen. Native and Se-Met BstC crystals appeared in the MCSG2-B3 condition containing 20% (v/v) PEG3500 and 0.2 M ammonium citrate dibasic after 2 weeks of incubation at 20 °C. Crystals were cryoprotected in the same conditions with the addition of 33% (v/v) PEG400 before being flash-cooled and stored in liquid nitrogen. Dimer-BstC crystals are obtained in the MCSG2-A12 condition containing 40 % (v/v) PEG 600 and 0.1 M Sodium Citrate: Citric Acid, pH 5.5 after 2 weeks of incubation at 20°C. Crystals were flash-cooled and stored in liquid nitrogen.

**Data collection, processing, and structure determination**

All the diffraction datasets were collected at 100 K at the Stanford Synchrotron Rdaiation Lightsource (SSRL, Stanford, CA, USA). BstB and dimer-BstC datasets were collected used a Pilatus 6M detector at beamline BL9-2, while BstC datasets were collected at beamline BL14-1 with a MARCCD325 detector. Native datasets were collected at a fixed wavelength at 0.97946 Å. MAD datasets were collected on ligand-free SeMet-substituted crystals at three wavelengths near the Se K-edge (0.97896 Å, 0.91837 Å, and 0.97930

Å). All diffraction datasets were indexed and processed by XDS (Kabsch, 2010a, 2010b) and HKL3000 packages (Minor et al., 2006) The structures of BstB and BstC were phased by the MAD method with Phenix AutoSol using the Se-anomalous diffraction (Liebschner et al., 2019). Subsequent density modification gave excellent electron-density maps, which allowed the building of the models. Iterations of refinement were carried out with Phenix-Refine, and model building was performed in Coot. The statistics of data collection and refinement are listed in Table S1.

All structural figures were prepared using PyMOL(Schrödinger and DeLano) or UCSF Chimera (Pettersen et al., 2004). Cavities in BstB were assessed using CASTp (Tian et al., 2018) with a probe radius of 1.4 Å, equivalent to the radius of water. The tunnel through BstC was identified with MOLE2.0 (Pravda et al., 2018) using default settings and a probe radius of 1.4 Å.

**Docking of substrate to BstB and BstC**

Molecular docking studies were performed using ICM-Pro 3.8-6a (Abagyan et al., 1994; Abagyan and Totrov, 1994). Briefly, chain A of the BstB and BstC crystal structure was used as the rigid receptor. Each protein molecule was converted to an ICM object, with optimization of hydrogen atom placement. Potential binding site were identified using the pocketfinder feature of ICM-Pro (Abagyan and Kufareva, 2009; An et al., 2005); for BstB and BstC the largest pockets had volumes of 1050 $A^3$ and 670 $A^3$, respectively. The coordinates of the C-4 methylated sterols (4-monomethylsterol and 4,4-dimethylsterol) were generated using the ICM-Pro ligand editing tools, based upon the structure of cholesterol. The ligands were docked into the binding pockets in both protein receptors. The MA-1-206 docking runs were performed multiple times, results were ranked in order to the overall score, and the most energetically favored binding modes were extracted from ICM-Pro as PDB files. Additional docking runs with BstB were carried out where three residues (Glu118, Tyr120 and Asn192) were allowed to have rotationally flexible side chains to simulate induced fit docking, using the explicit group docking feature of ICM-Pro, while the rest of the protein remained rigid. Result analyses and figure rendering were performed using PyMOL (Schrödinger and DeLano).

**Molecular dynamics simulations of BstB and BstC**

Molecular dynamics (MD) simulations were performed in triplicate on models comprising BstB and BstC docked with 4-monomethylsterol and 4,4-dimethylsterol, using Desmond (Bowers et al., 2006) in the Schrodinger 2019-2 release. The docked complexes were prepared with Maestro (Schrodinger) using the OPLS3e force field (Roos et al., 2019). The pre-defined TIP3P water model (Neria et al., 1996) was used to build the system. The overall charge of the complexes was calculated as -1 and -10 for BstB and BstC respectively and neutralized with $Na^+$ ions. Prior to building in system for both complexes, 0.15 M salt (NaCl) was added, and the ions were restricted from coming within 20 Å of the ligand. The systems were minimized prior to the final 20 ns production step runs at 300 K and 1 atm pressure, using the Nosé–Hoover chain coupling scheme for the temperature control and the Martyna–Tuckerman–Klein chain coupling scheme with a coupling constant of 2.0 ps for pressure control (Martyna et al., 1992). Nonbonded forces were calculated using an r-RESPA integrator. The trajectories were saved at 10 ps intervals for analysis. For the 150 ns simulation on apo-BstB, the same protein preparation steps and variable settings were used except the trajectories were saved every 25 ps for analysis. Maestro and Desmond were run on the SHERLOCK 3.0 HPC cluster at Stanford University.

### Supporting figures and legends

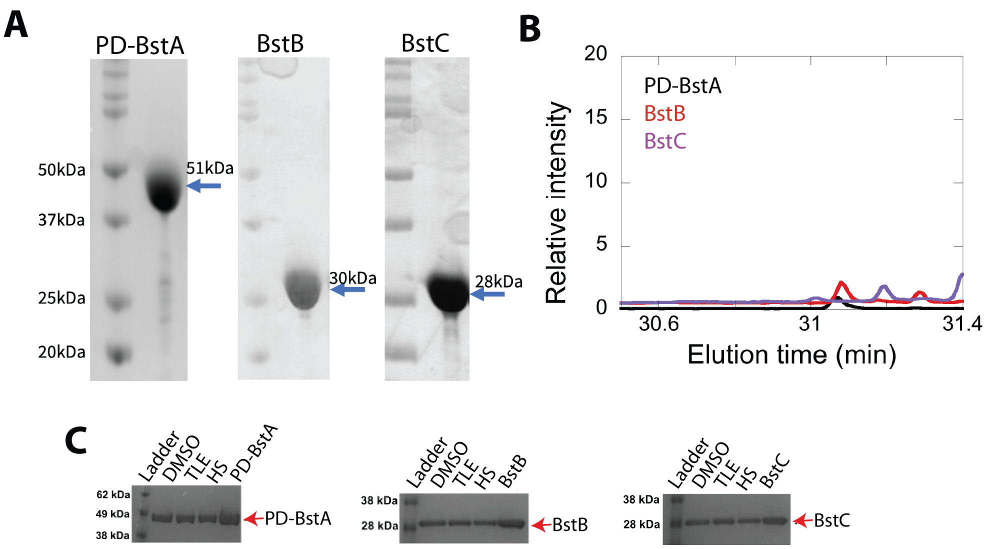


**Figure S1.** (**A**) SDS-PAGE gel of pure $BstA^{PD}$, BstB, and BstC. All proteins are estimated to be more than 90% pure. (**B**) GC chromatograms of protein-lipid pull down control experiments (DMSO-treated samples) reveal that the proteins do not contain C4-methylsterols when *M. capsulatus* lipids (TLE or HS fraction) are not present. **(C)** SDS PAGE gels demonstrating that the samples analyzed in the protein-lipid pull down assays did contain $BstA^{PD}$, BstB, and BstC.

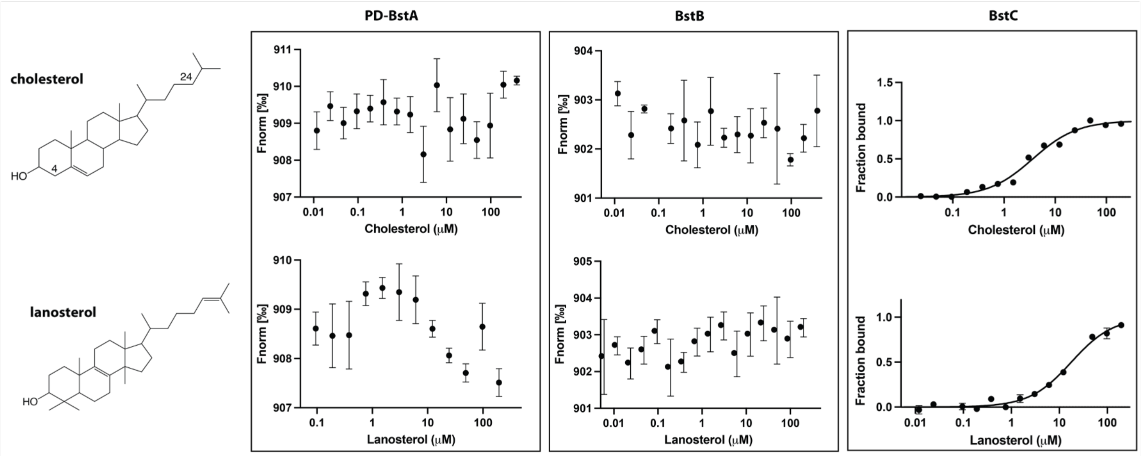


**Figure S2.** Binding of cholesterol (top) and lanosterol (bottom) to BstA$^{PD}$, BstB, and BstC. The data were acquired using the same conditions as reported for **Figure 2.** Each measurement was performed 3 times, but the average value reported as points and the standard deviation shown as error bars.

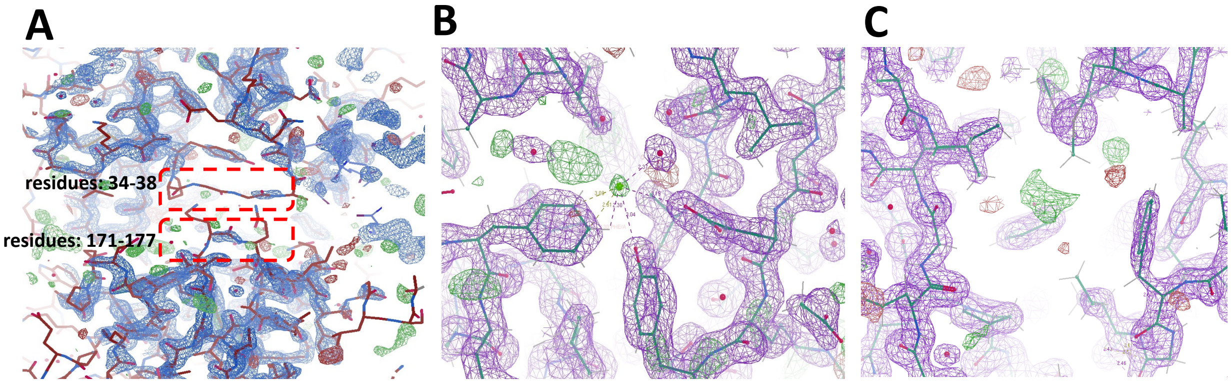


**Figure S3.** A portion of the 2Fo − Fc electron density omit map contoured at 2 σ and the Fo−Fc electron density omit map contoured at 3σ (green mesh) of BstB. **(A)** Loops (residues 34-38 and 171-177) are shown in red boxes. The poor density of these loops could be due to protein flexibility. **(B)** The modeled chloride forms polar interactions with the surrounding Glu118, Tyr120 and Phe150. **(C)** Unmodeled density found in the cavity where the aliphatic tail of the sterol is positioned in the docking model.

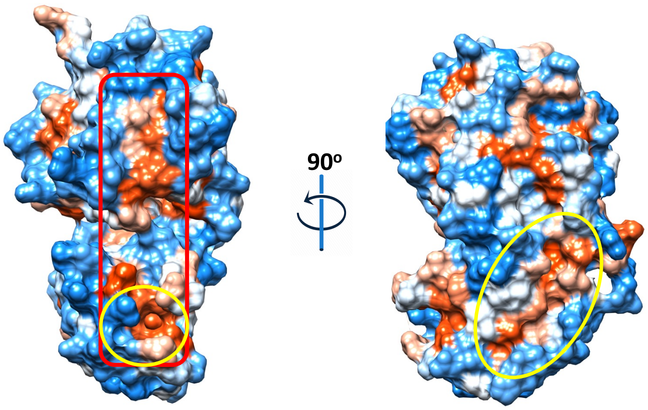


**Figure S4.** Hydrophobicity representation of the BstB structure shows a second hydrophobic pocket located in domain B below the missing loop (202-204). The pocket (circled) has an area of 172 $Å^2$ and a volume of 117 $Å^3$ and may function as a second substrate binding site. The surface in red box is the potential interface after substrate binding.

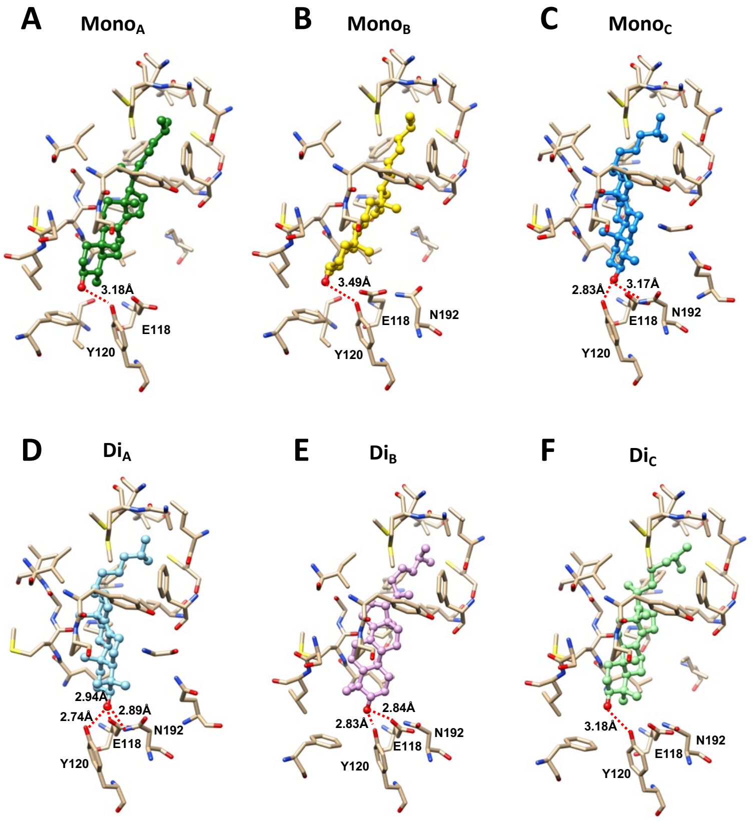


**Figure S5.** Docking models of 4-monomethyl sterol and 4,4-dimethyl sterol in BstB structure. **(A-C)** Energetically favored poses of 4-monomethylsterol. **(D-F)** Energetically favored poses of 4,4-dimethylsterol. Sterols are shown in ball and stick with the hydroxyl group is colored in red; the surrounding residues are shown in sticks and colored in tan. Hydrogen bonds between the hydroxyl group of the sterol and surrounding residues are shown as red dashed lines.

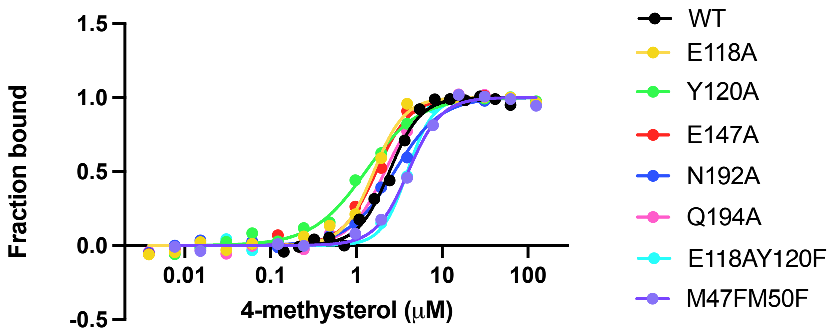


| | WT | E118A | Y120A | E147A | N192A | Q194A | E118A /Y120F | M47F /M50F |
|---|---|---|---|---|---|---|---|---|
| $n_H$ | 2.4 | 2.4 | 1.4 | 2.2 | 1.6 | 2.1 | 2.9 | 2.3 |
| $K_{d\ avg}$ (μM) | 2.4 | 1.6 | 1.3 | 1.7 | 2.8 | 2.1 | 3.9 | 4 |

**Figure S6**. MST binding curves of BstB variants with 4-monomethyl sterol.

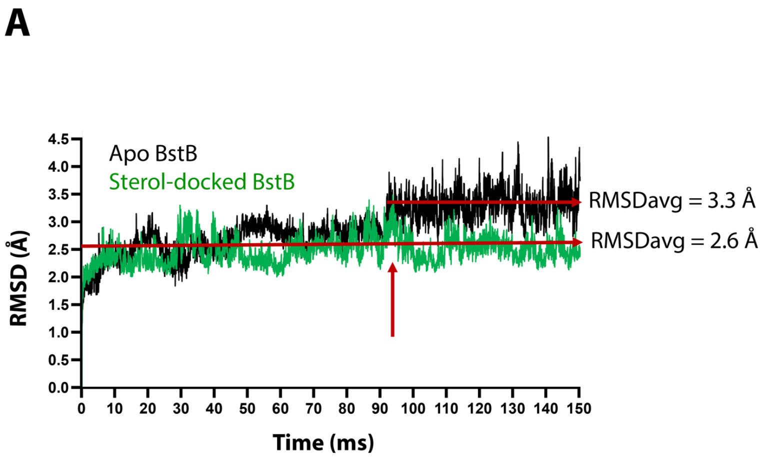


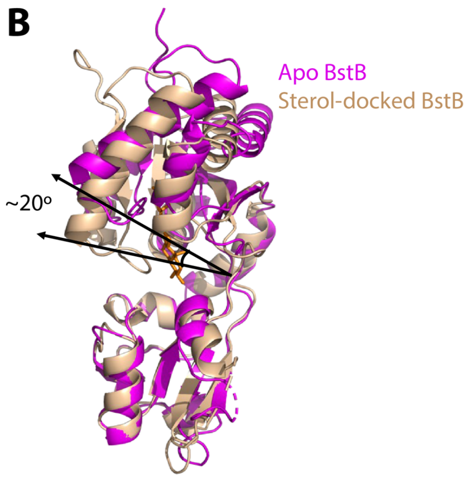


**Figure S7. (A)** RMSD vs time plot for the MD simulation on the apo-BstB and docking complex with 4-monomethyl sterol. The trajectory of apo-BstB is in black while that of the docking complex is depicted in green. **(B)** Superposition of the apo-BstB structure with the monomethyl sterol-docked complex.

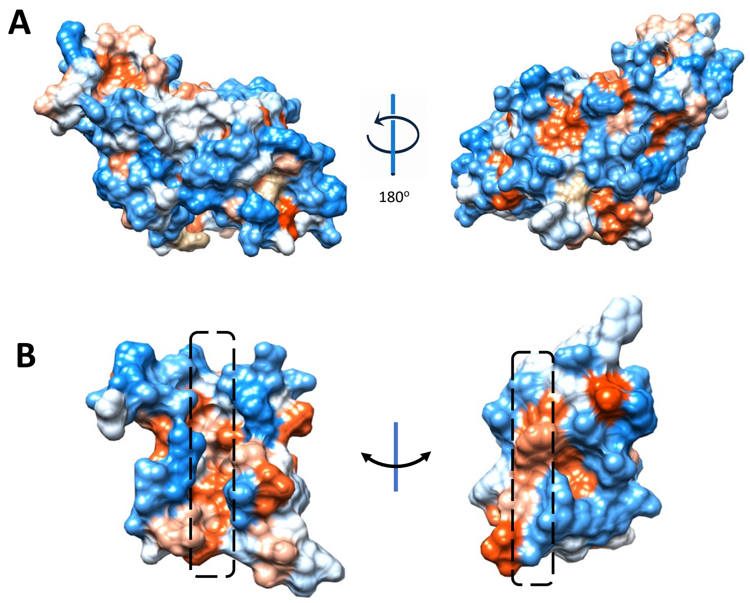


**Figure S8**. Hydrophobicity representations of the BstC structure on **(A)** the exterior surface and **(B)** the cross-section of the side view of the tunnel in BstC structure. TN1 exhibits hybrid hydrophobicity, with the sides horizontal to cavities being hydrophobic while the vertical sides are more hydrophilic.

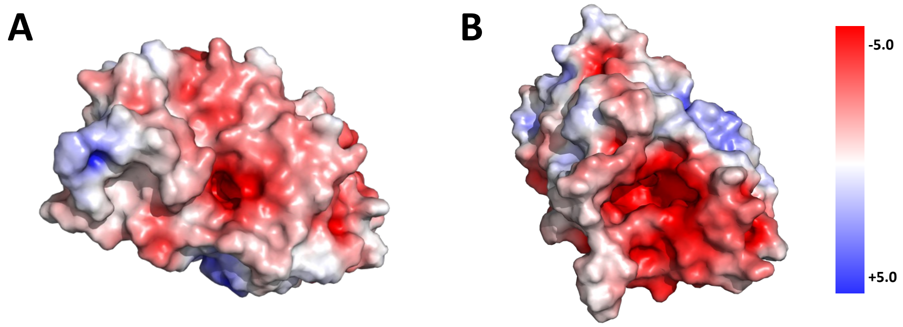


**Figure S9**. Electrostatic calculation shows that most of the surface, including the N-terminal **(A)** and C-terminal **(B)** terminal faces, exhibits strong negative charge.

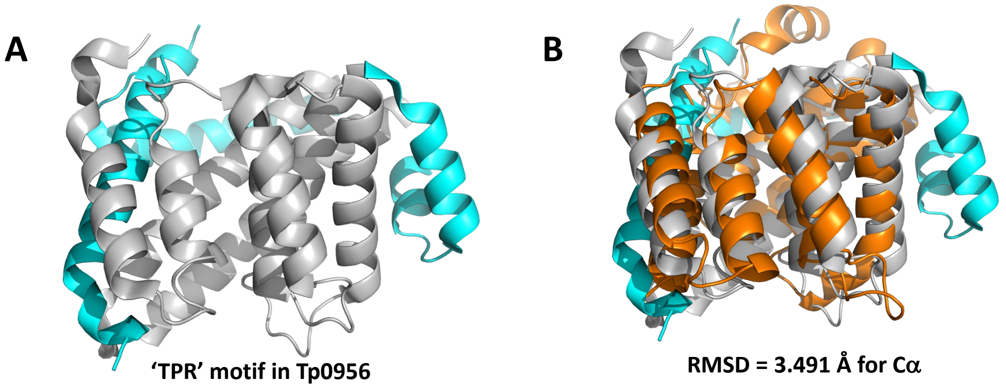


**Figure S10.** (**A**) Cartoon depiction of Tp0956 (PDB 3U64), the structural homolog of BstC. The protein contains four helical hairpins that are similar to tetratricopeptide repeat (TPR) motifs, which are hallmarks of proteins that engage in protein–protein interactions. The TPR motif was colored in grey. (**B**) Superposition of the BstC structure with Tp0956 (RMSD of 3.491 Å for all Cαs). BstC is colored in orange.

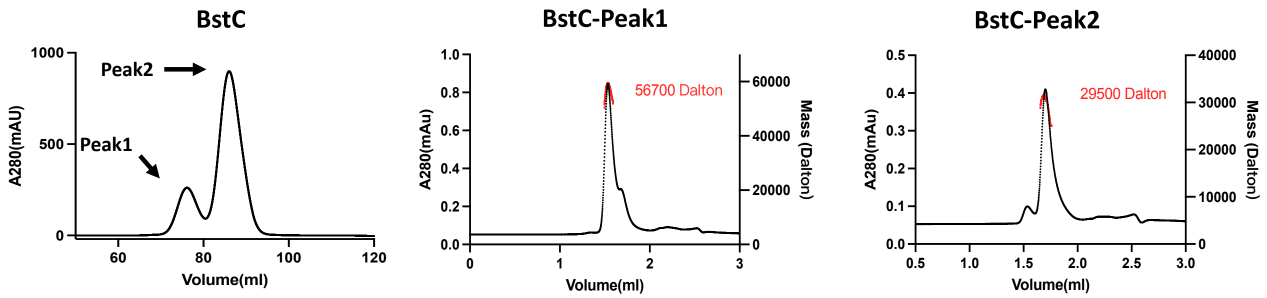


**Figure S11.** Size exclusion chromatography (SEC, left) and SEC-multiangle light scattering (MALS, middle and right) analysis of BstC. Two peaks are observed in the SEC chromatogram, with SEC-MALS revealing that they correspond to monomeric and dimeric BstC.

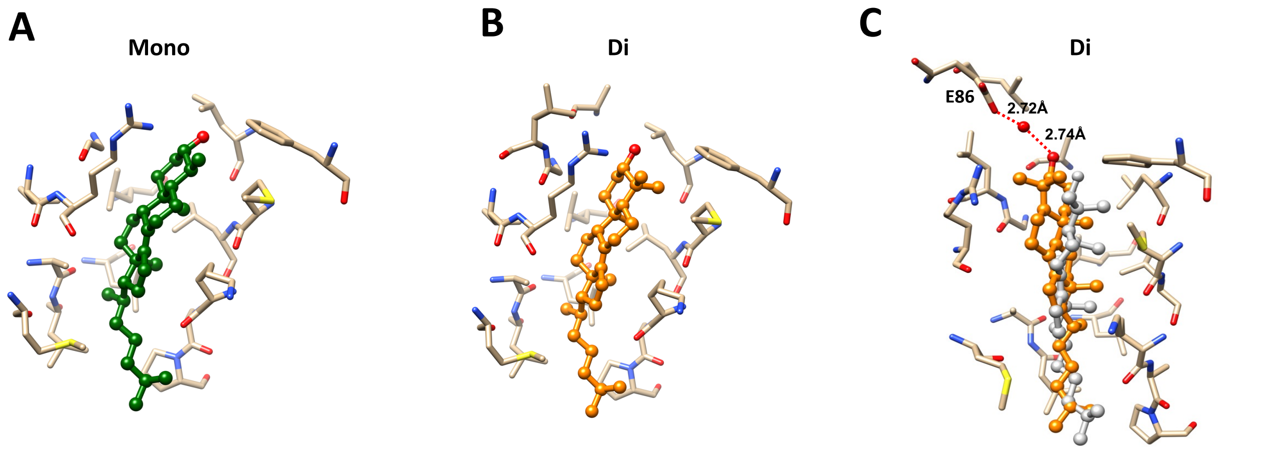


**Figure S12**. Docking models of 4-monomethyl sterol and 4,4-dimethyl sterol in BstC structure. **(A)** Energetically favored poses of 4-monomethyl sterol. The sterol is colored in green. (**B)** Energetically favored poses of 4,4-dimethylsterol. The sterol is colored in orange **(C)** Docking complex of 4,4-dimethylsterol before and after MD simulation. The sterol prior to the MD simulation is colored in grey and while orange shows its position after the simulation. Sterols are shown in ball and stick with the hydroxyl group is colored in red; the surrounding residues are shown in sticks and colored in tan. Water-mediated hydrogen bonding between the hydroxyl group of the sterol and E86 is highlighted.

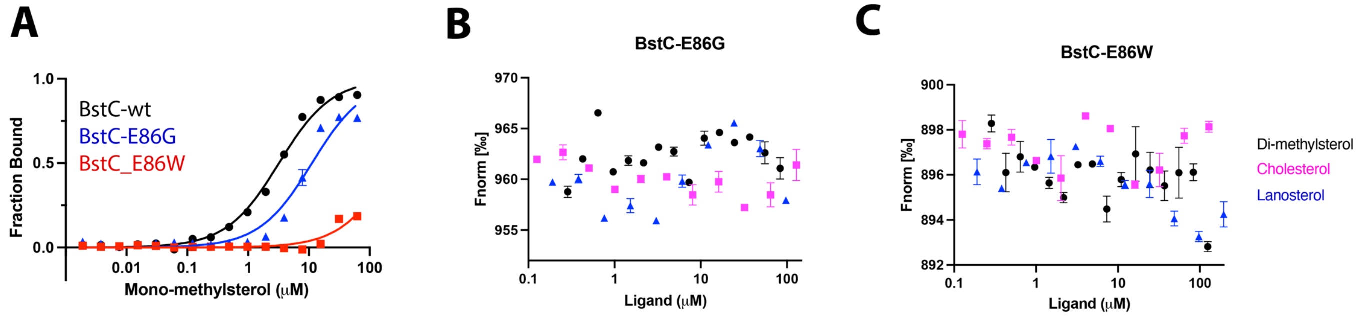


**Figure S13**. MST analysis of the BstC variants (50 nM) binding to sterols. **(A)** The E86G and E86W variants bind to C4-mono-methyl sterol with $K_d$s of 11.2±2.93 μM and 250.4±538.4 μ **(B,C)** No interaction was detected between BstC E86 variants and other sterols.

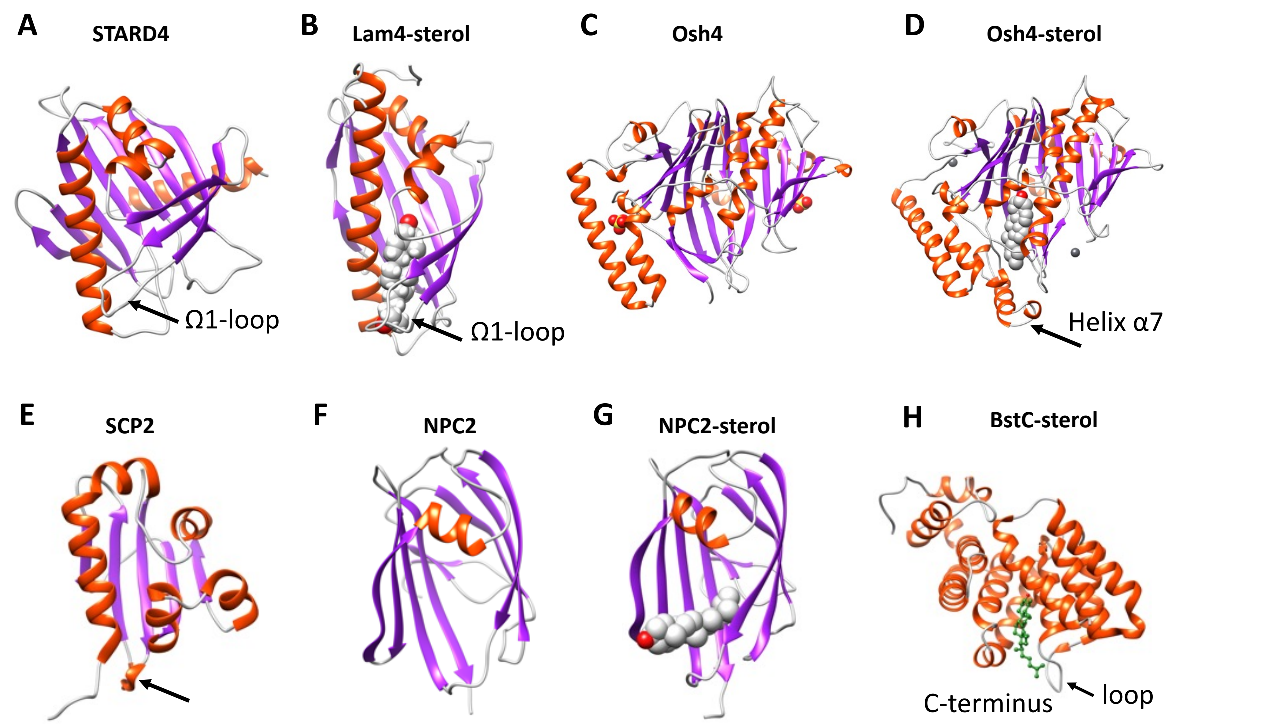


**Figure S14**. Structural comparison of eukaryotic sterol-binding proteins.
**(A)** Apo human STARD4 (PDB 6L1D). **(B)** Yeast Lam4 bound to 25-hydroxycholesterol (PDB 6BYM). The Ω1-loop is depicted in structures that possess it. **(C-D)** Yeast apo- (**C**, PDB 1ZI7) and cholesterol-bound (**D**, PDB 1ZHY) Osh4. The N-terminal helix α7 in Osh4, which is missing in the apo-Osh4 structure, may work similar to the Ω1-loop. **(E)** The non-specific LTP, rabbit SCP2 (PDB 1C44); **(F)** apo- (PDB 6R4M) and (**G**) ergosterol-bound (PDB 6R4N) NPC2. All structures are shown in cartoon representation with the α-helixes and β-strands colored in red and magenta, respectively. The sterol substrates are depicted in grey spheres with the hydroxyl group(s) colored in red. **(H)** The Ω1-like loop in BstC docked complex with monomethyl sterol.

**Table S1. Available structural information in the SSN of different pfam members.**

| Superfamily | Pfam | Protein | PDB | Organism |
|---|---|---|---|---|
| MMPL | PF03176 | HpnN | 5khs | *Burkholderia multivorans* |
| MMPL | PF03176 | MmpL3 | 6ajf | *Mycobacterium smegmatis* |
| MMPL | PF03176 | MmpL11 | 4y0l | *Mycobacterium tuberculosis* |
| Phosphonate-bd | PF12974 | AioX | 6esk | *Pseudorhizobium banfieldiae* |
| Phosphonate-bd | PF12974 | PhnD | 5lq5 | *Prochlorococcus marinus* |
| Phosphonate-bd | PF12974 | PtxB | 5lv1 | *Prochlorococcus marinus* |
| Phosphonate-bd | PF12974 | PtxB | 5o2k | *Pseudomonas stutzeri* |
| Phosphonate-bd | PF12974 | PtxB | 5jvb | *Trichodesmium erythraeum* |
| Phosphonate-bd | PF12974 | PA3383 | 3n5l | *Pseudomonas aeruginosa* |
| Phosphonate-bd | PF12974 | PhnD | 3qk6 | *Escherichia coli* |
| Phosphonate-bd | PF12974 | XAC2383 | 5ub3 | *Xanthomonas axonopodis pv. citri* |
| Phosphonate-bd | PF12974 | HtxB | 5me4 | *Pseudomonas stutzeri* |
| TatT | PF16811 | Tp0956 | 3u64 | *Treponema pallidum (strain Nichols)* |

**Table S2. Data collection and refinement statistics.**

| | apo-BstB(SeMet) | apo-BstC(SeMet) |
|---|---|---|
| **Resolution range** | 38.4 -1.6 (1.657-1.6) | 28.63 -1.91 (1.978-1.91) |
| **Space group** | P 21 21 21 | P 1 21 1 |
| **Unit cell (a, b, c; α, β, γ)** | 39.882, 40.355, 142.263; 90, 90, 90 | 55.083, 57.257, 85.825; 90, 97.26, 90 |
| **Total reflections** | 606476 (60185) | 200826 (19731) |
| **Unique reflections** | 31071 (3061) | 41329 (4084) |
| **Multiplicity** | 19.5 (19.7) | 4.9 (4.8) |
| **Completeness (%)** | 97.81 (93.74) | 99.91 (99.98) |
| **Mean I/sigma(I)** | 30.96 (2.07) | 11.09 (2.78) |
| **Wilson B-factor** | 22.86 | 15.59 |
| **R-merge** | 0.07677 (1.462) | 0.1267 (0.6131) |
| **R-meas** | 0.07881 (1.501) | 0.1423 (0.6891) |
| **R-pim** | 0.01755 (0.3345) | 0.06419 (0.3117) |
| **CC1/2** | 1 (0.745) | 0.994 (0.762) |
| **CC*** | 1 (0.924) | 0.999 (0.93) |
| **Reflections used in refinement** | 30554 (2873) | 41317 (4084) |
| **Reflections used for R-free** | 1964 (184) | 1996 (198) |
| **R-work** | 0.2038 (0.2892) | 0.174(0.2389) |
| **R-free** | 0.2234 (0.2749) | 0.207 (0.2675) |
| **CC(work)** | 0.955 (0.786) | 0.948 (0.879) |
| **CC(free)** | 0.948 (0.725) | 0.939 (0.834) |
| **Number of non-hydrogen atoms** | 2117 | 4036 |
| **macromolecules** | 1974 | 3546 |
| **solvent** | 142 | 355 |
| **Protein residues** | 257 | 464 |
| **RMS(bonds)** | 0.010 | 0.012 |
| **RMS(angles)** | 1.03 | 1.28 |
| **Ramachandran favored (%)** | 99.21 | 100.00 |
| **Ramachandran allowed (%)** | 0.79 | 0.00 |
| **Ramachandran outliers (%)** | 0.00 | 0.00 |
| **Rotamer outliers (%)** | 0.97 | 0.00 |
| **Clashscore** | 2.28 | 1.39 |
| **Average B-factor** | 33.82 | 19.33 |
| **macromolecules** | 33.84 | 18.22 |
| **ligands** | 30.00 | 27.28 |
| **solvent** | 33.49 | 27.44 |
| **Number of TLS groups** | 3 | 1 |

Statistics for the highest-resolution shell are shown in parentheses.

**Table S3.** Docking and molecular dynamics results for BstB.

| | | ICM docking results energies (kcal/mol)[a] | | | | Hydrogen bonds in docked poses distance (Å) | | | Hydrogen bonds after MD bond persistence (%) | | |
|---|---|---|---|---|---|---|---|---|---|---|---|
| **Ligand** | **Pose** | **Score[b]** | **Hbond** | **Hphob** | **VwInt** | **Glu118** | **Tyr120** | **Asn192** | **Glu118** | **Tyr120** | **Asn192** |
| Mono | A | -17.7 | -0.94 | -9.62 | -25.60 | - | 3.18 | - | 95 | 3 | - |
| | B | -20.4 | -0.69 | -9.50 | -26.10 | - | 3.49[c] | - | 95 | 2 | 1 |
| | C | -16.9 | -2.77 | -9.32 | -23.73 | - | 2.83 | 3.17 | 85 | - | 11 |
| Di | A | -20.3 | -4.53 | -9.48 | -23.22 | 2.94 | 2.74 | 2.89 | 12 | 12 | 57 |
| | B | -27.8 | -4.64 | -9.74 | -28.74 | 2.84 | 2.83 | - | 94 | 10 | 1 |
| | C | -17.2 | -1.99 | -9.94 | -24.13 | - | 3.18 | - | - | -[d] | - |

[a] Energies for hydrogen bonding (Hbond), hydrophobic interactions (Hphob) and van der Waals interactions (VwInt).
[b] The score has no units. More negative numbers are indicative of stronger binding.
[c] This distance could be considered too long to be a hydrogen bond.
[d] This pose loses the hydrogen bond with Tyr120 but gains a hydrogen bond with the carbonyl oxygen of Pro34 which then persists for 95% of the MD simulation.

**Table S4. Residues involved in forming cavity in cleft of BstB.**

| Domain | SeqID | AA | Domain | SeqID | AA |
|---|---|---|---|---|---|
| A | 29 | Val | A | 115 | Arg |
| A | 31 | Val | A | 117 | Ser |
| A | 33 | Tyr | A | 118 | Glu |
| A | 34 | Pro | B | 120 | Tyr |
| A | 35 | Gly | B | 143 | Thr |
| A | 36 | Gly | B | 144 | Met |
| A | 37 | Ala | B | 146 | Asp |
| A | 38 | Val | B | 147 | Glu |
| A | 39 | Asn | B | 150 | Phe |
| A | 40 | Glu | B | 171 | Ser |
| A | 43 | Ala | B | 172 | Arg |
| A | 44 | Asp | B | 173 | Gln |
| A | 46 | Ala | B | 174 | Ala |
| A | 47 | Met | B | 175 | Ile |
| A | 48 | Asp | B | 192 | Asn |
| A | 49 | Ala | B | 193 | Glu |
| A | 50 | Met | B | 194 | Gln |
| A | 51 | Leu | B | 195 | Gln |
| A | 53 | Val | B | 217 | Ile |
| A | 54 | Val | A | 218 | Pro |
| A | 67 | Ser | A | 219 | Leu |
| A | 69 | Phe | A | 221 | Gly |
| A | 71 | Ala | A | 222 | Val |
| A | 73 | Val | A | 223 | Val |

| | | | | | |
|---|---|---|---|---|---|
| A | 89 | Ile | A | 237 | phe |
| A | 90 | Thr | A | 241 | Leu |
| A | 91 | Ser | A | 244 | Leu |
| A | 92 | Leu | A | 253 | Leu |
| A | 93 | Ala | A | 254 | Cys |
| A | 94 | Leu | A | 256 | Leu |
| A | 95 | Tyr | A | 257 | Phe |
| A | 106 | Pro | A | 258 | Gly |
| A | 108 | Val | A | 259 | Ile |
| A | 109 | Gln | A | 262 | Phe |
| A | 110 | Pro | A | 270 | Phe |
| A | 112 | Ile | | | |

### Table S5. Residues involved in forming cavity in BstC.

| Tunnel1 | Chamber2 | Chamber3 |
|---|---|---|
| ASP 45 | GLY 128 | LEU 129 |
| PRO 49 | LEU 129 | ARG 132 |
| LEU 82 | ARG 132 | LEU 167 |
| HIS 83 | MET 135 | MSE 170 |
| GLU 86 | ALA 138 | LYS 174 |
| TYR 121 | LEU 139 | LEU 207 |
| TYR 122 | LEU 142 | PHE 208 |
| LEU 129 | LEU 145 | GLU 211 |
| ARG 132 | LEU 167 | GLU 251 |
| ASP 160 | MSE 170 | ASP 254 |
| ARG 166 | LEU 171 | LEU 255 |
| LEU 167 | LYS 174 | |
| LEU 204 | PHE 208 | |
| TRP 241 | | |
| ASN 244 | | |
| TRP 248 | | |

### Table. S6 Protein sequences and primers.

| NAME | SEQUENCE | PRIMERS |
|---|---|---|
| **BstA** | MNIPHLAALAAERFARRPWRVLALAMALSALSLWAVSRL PVHTSRQALLPHDNAVAQRFDAFLDKFGAASDLIVVLEG APPDELKPFADELATALAAEPEIAQATARLDLRFVLEHAY LAVTPERLGTLAGVLEKFGAGAIPEDSSQVDATLGRLLQW LEGAPAMPAAGIDLPTVEVGLKLLGASLDEWHRWLSAGE VPAALDWTRLLAGLGGSEIANDGYFVSRDGRMYFLFVHP ASASEDFTAIGPFVEKVRTVAADRAARARAAGRTAPKVG LTGLPAIEYEEHVSIRHDIALVVGSAAGLIVLLILVVVRSW RWALVIFVPMGLGVLWSLGLALVTIGHLTLITASFIAVLFG LGADYGIFTSARIAEERRRGKPLTEAIGAGMGASFQAVFT AGGASVVIFGALATVDFPGFSELGLVAAKGVMLILVSTWL VQPALYALLPPKLAPLPAAASAGAIEPGRMPFRGSVAVIL VAGALATAAFGIGSGYELPFDYDVLSLLPKDSESAYYQNR MVAESDYQAEVVIFTAPDLEEARRIAAEAGRLGSVAKVQ | pET20b-Forward: GAAGGAGATATACATATGAACATCCCC CATTTGGCCGCTC<br>pET20b-Reverse: GTGGTGGTGGTGCTCGAGTGGTTTCTTT CTTGAATCGAGAAGC<br>pET28a-Forward: AGAATCTTTATTTTCAGGGCCATAACAT TCCGCACCTGGCGGC<br>pET28a-Reverse: CTCAGTGGTGGTGGTGGTGGTGCTCGAG CTTAGGTTTCTTACGGCTGTCCAG |

| | | |
|---|---|---|
| | SLMDLFPPDADARALEARRIGELADDGYAVRLARLAAIGL<br>PEGTFGRVRTILEKGGDFIDQSQELAFSAGHSGLVAALED<br>VRGRLDAVRSAIEADPVQARERSERFFRMLLSAAERGVAL<br>LAEWRQARPITPAQLPPALRDRFFAADGTVAVYAFPAKT<br>VYDPANLDRLMQEIYGVSPDATGFPATHQVFSKSVVESFT<br>HGTREAVTVCLLWLALVLRNWRGFVLASMPLLIGGGWM<br>LGLMALCGIRYNYANIIALPLVIALAVDYGVWFSQRWFDL<br>KDRSLTQINRVAGGVIGLAAGTELAGLGAITLANYRGVSS<br>LGVNITVGLLCCLAATLWVAPAIGQLLDSRKKP | |
| **BstA$^{PD}$** | MAVAQRFDAFLDKFGAASDLIVVLEGAPPDELKPFADEL<br>ATALAAEPEIAQATARLDLRFVLEHAYLAVTPERLGTLAG<br>VLEKFGAGAIPEDSSQVDATLGRLLQWLEGAPAMPAAGI<br>DLPTVEVGLKLLGASLDEWHRWLSAGEVPAALDWTRLL<br>AGLGGSEIANDGYFVSRDGRMYFLFVHPASASEDFTAIGP<br>FVEKVRTVAADRAARARAAGRTAPKVGLT**GSGSGSGS**ES<br>AYYQNRMVAESDYQAEVVIFTAPDLEEARRIAAEAGRLG<br>SVAKVQSLMDLFPPDADARALEARRIGELADDGYAVRLA<br>RLAAIGLPEGTFGRVRTILEKGGDFIDQSQELAFSAGHSGL<br>VAALEDVRGRLDAVRSAIEADPVQARERSERFFRMLLSA<br>AERGVALLAEWRQARPITPAQLPPALRDRFFAADGTVAV<br>YAFPAKTVYDPANLDRLMQEIYGV | pET28a-Forward1:<br>CTTTATTTTCAGGGCCATATGGCGGTGG<br>CGCAACGTTTCGAC<br>pET28a-Reverse1:<br>AGTACGCGCTCTCGCTACCGCTACCGCT<br>GCCGCTACCGGTCAGACCAACTTTCGGC<br>pET28a-Forward2:<br>GTTGGTCTGACCGGTAGCGGCAGCGGT<br>AGCGGTAGCGAGAGCGCGTACTATCAG<br>AAC<br>pET28a-Reverse2:<br>CTCAGTGGTGGTGGTGGTGGTGCTCGAG<br>TCACACACCATAAATTTCCTG |
| **BstB** | M**TDKITCFSLLAALLAPFVPAQA**GAPAPVVVCYPGGAV<br>NERDADQAMDAMLRVVERVGQWPEKSFSSVFTAKVADC<br>GKLMAEMKPAFAITSLALYLDMRGQYDLVPVVQPRIDGR<br>TSERYRVVAKQGRFHDMDELKGRTLGGTMLDEPAFLGKI<br>VFAGKYDPEKDFALQPSRQAIRALRSLDKGELDAVVLNE<br>QQFAGLSALQLASPVETVFTSAEIPLMGVVANARLTSAQE<br>RARFAQALETLCADPEGRKLCDLFGIQSFVAVDPTVFDPM<br>ARLWLARN | pET20b-Forward:<br>GAAGGAGATATACATATGGGCGCCCCC<br>GCACCG<br>pET20b-Reverse:<br>GTGGTGGTGGTGCTCGAGGTTCCGCGCC<br>AGCCA<br>pET28a-Forward:<br>CGAGAATCTTTATTTTCAGGGCCATGGC<br>GCCCCCGCACCGGTGGTC<br>pET28a-Reverse:<br>CTCAGTGGTGGTGGTGGTGGTGCTCGAG<br>TTAGTTCCGCGCCAGCCAGAG |
| **BstC** | M**RLFAAGILAGVLAG**CGGLHRDGTPAGPSAGCPRLTAA<br>ALSAGQDALGPSSETQELECALDFLRGSDDPALRRSSLGS<br>RICLHLAERNSDPAERARFAREGVERAEAALAQGGEDDG<br>AVHYYLAANLGLAVRDDMTAALANLHRLEHESEAAVKL<br>SPDFDDGGPLRLLGMLYLKAPAWPAGMGDGDKALDLLG<br>QAVERHPGHPLNHLFYAEALWEVNGESESRRVEEEMAAG<br>WRLLESGSWGYNKQIWKREFADLRQEIGAPAR | pET20b-Forward:<br>GAAGGAGATATACATATGTGCGGCGGT<br>CTTCAC<br>pET20b-Reverse:<br>GTGGTGGTGGTGCTCGAGCCTAGCGGG<br>CGCCCC<br>pET28a-Forward:<br>ATCTTTATTTTCAGGGCCATTGCGGCGG<br>TCTTCACCGCGATG<br>pET28a-Reverse:<br>CTCAGTGGTGGTGGTGGTGGTGCTCGAG<br>TTACCTAGCGGGCGCCCCGAT |

* The signal peptides in BstB and BstC are shown in bold.

**Movie 1. 150 ns molecular dynamics (MD) simulations on apo-BstB structure.**

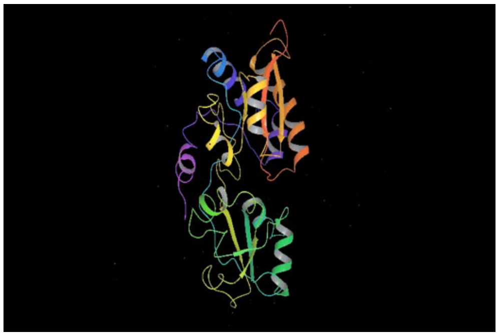

**Movie 2. 150 ns molecular dynamics (MD) simulations on *Mono$_C$*-docked BstB complex.**

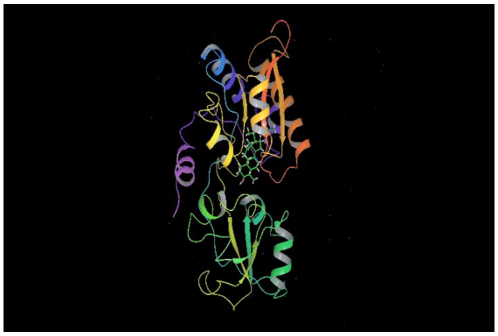
